# scRNA-seq reveals transcriptional dynamics of *Encephalitozoon intestinalis* parasites in human macrophages

**DOI:** 10.1101/2024.05.30.596468

**Authors:** Pattana Jaroenlak, Kacie L. McCarty, Bo Xia, Cherry Lam, Erin E. Zwack, Itai Yanai, Gira Bhabha, Damian C. Ekiert

## Abstract

Microsporidia are single-celled intracellular parasites that cause opportunistic diseases in humans. *Encephalitozoon intestinalis* is a prevalent human-infecting species that invades the small intestine. Dissemination to other organ systems is also observed, and is potentially facilitated by macrophages. The macrophage response to infection and the developmental trajectory of the parasite are not well studied. Here we use single cell RNA sequencing to investigate transcriptional changes in both the host and parasite during infection. While a small population of infected macrophages mount a response, most remain transcriptionally unchanged, suggesting that the majority of parasites may avoid host detection. The parasite transcriptome reveals large transcriptional changes throughout the life cycle, providing a blueprint for parasite development. The stealthy microsporidian lifestyle likely allows these parasites to harness macrophages for replication and dissemination. Together, our data provide insights into the host response in primary human macrophages and the *E. intestinalis* developmental program.

## INTRODUCTION

Microsporidia are obligate, intracellular parasites closely related to fungi^1^. Over 1,500 species have been reported, which infect a wide range of invertebrate and vertebrate hosts, including humans^2^. Microsporidia exist in the environment as dormant spores surrounded by a thick spore coat. In humans, transmission of microsporidia primarily occurs through ingestion of contaminated food or water. Consequently, the initial site of infection is frequently the gastrointestinal tract^3^. Microsporidia infection often results in self-limiting diarrheal disease in otherwise healthy individuals. However, in immunocompromised patients, infection can be fatal^4^, and disseminated disease has been reported to cause encephalitis^5^, tracheobronchitis^6^, nephritis^7^, keratoconjunctivitis^8^, and myositis^9^.

*Encephalitozoon intestinalis* is one of the most common microsporidian species found to infect humans. The initial site of infection is usually the small intestine^10^, and in some cases the infection can spread systemically^11^. Dissemination is thought to occur via migrating macrophages, which can act as vehicles for parasites to spread throughout the body^12^. *E. intestinalis* is known to infect macrophages in culture^13,14^, as well as in the intestinal lamina propria of human patients^15^. These infection events have been proposed to be important for both parasite replication and dissemination within the host. Macrophages are enriched in the intestinal lamina propria and are among the first responders to incoming infection. Like other intracellular pathogens, microsporidia have developed mechanisms to evade killing by macrophages. Studies have shown that *Encephalitozoon* spp. are able to establish infection within macrophages by preventing phagosome acidification and inhibiting fusion with lysosomes^16^. Furthermore, *Encephalitozoon* spp. were also observed to suppress apoptosis in human macrophages, which prevents the killing of the parasites^17^.

To understand how *E. intestinalis* manipulates human macrophages to promote replication and spread, it is necessary to understand the transcriptional dynamics of how macrophages respond to infection, as well as the developmental program of the parasites. Previous work using bulk RNA sequencing of *E. intestinalis*-infected Caco-2 cells, a human colon cancer cell line, revealed that infection leads to mitochondrial stress and has an impact on cell signaling networks related to energy, metabolism, and membrane trafficking^18^. While bulk RNA sequencing can provide insights into the host-parasite interface across the whole population, it does not capture the complexity and heterogeneity of infection dynamics at the cellular level. Furthermore, no transcriptomic studies have been performed on other infected cell types. To understand how asynchronously replicating *E. intestinalis* parasites propagate within macrophages and evade detection, we performed single cell RNA sequencing (scRNA-seq) of primary human macrophages infected with *E. intestinalis* over their life cycle. From these data we analyzed both the host and parasite transcriptomes. Our results reveal that most macrophages fail to respond to invading parasites and are transcriptionally indistinguishable from uninfected cells, suggesting that *E. intestinalis* generally avoids detection, enabling successful parasite replication. In addition, we generate a transcriptional blueprint for parasite development, facilitating the identification of molecular markers at each stage of parasite development.

## RESULTS

### *E. intestinalis* infection dynamics in primary human macrophages

To identify suitable timepoints for scRNA-seq analysis, we isolated monocyte-derived macrophages from human donors and investigated the timeline of the *E. intestinalis* life cycle in these cells. We infected macrophages with purified *E. intestinalis* spores, and monitored infection by fluorescence microscopy from 3 hours post infection (hpi) to 72 hpi (Fig. 1a-e). At each timepoint, infected cultures were co-stained for DNA (DRAQ5) and chitin (calcofluor white), allowing for the identification of mature spores (DNA puncta surrounded by a chitin coat), as well as actively replicating parasites (DNA puncta lacking a chitin coat, inside macrophages). At 3 hpi and 12 hpi, we were unable to identify any replicating parasites. However, many macrophages contain ∼1-5 mature *E. intestinalis* spores at these early time points (Fig. 1a,b). As productive infection typically yields dozens of spores and takes ∼48 hrs, we infer that macrophages containing ≤5 mature spores and no replicating parasites are likely the result of phagocytosis of spores from the inoculum (Fig. 1a,b). By 24 hpi, we observed clusters of replicating parasites in ∼5% of macrophages (Fig. 1c,f), which is earlier than previously reported^12^. Consistent with previous infection models for *E. intestinalis*^19^, replicating parasites are clustered, likely within the parasitophorous vacuole (Fig. 1c). By 48-72 hpi, ∼20-30% of the macrophages contain newly replicated parasites (Fig. 1d-f). Many of these parasites are actively replicating, while others have matured into sporoblast or mature spore stages, as evidenced by the acquisition of a chitin coat. Together, our data suggest that replicating parasites are most abundant 24-48 hpi, with spore maturation occurring ∼48-72 hpi.

**Figure 1.**
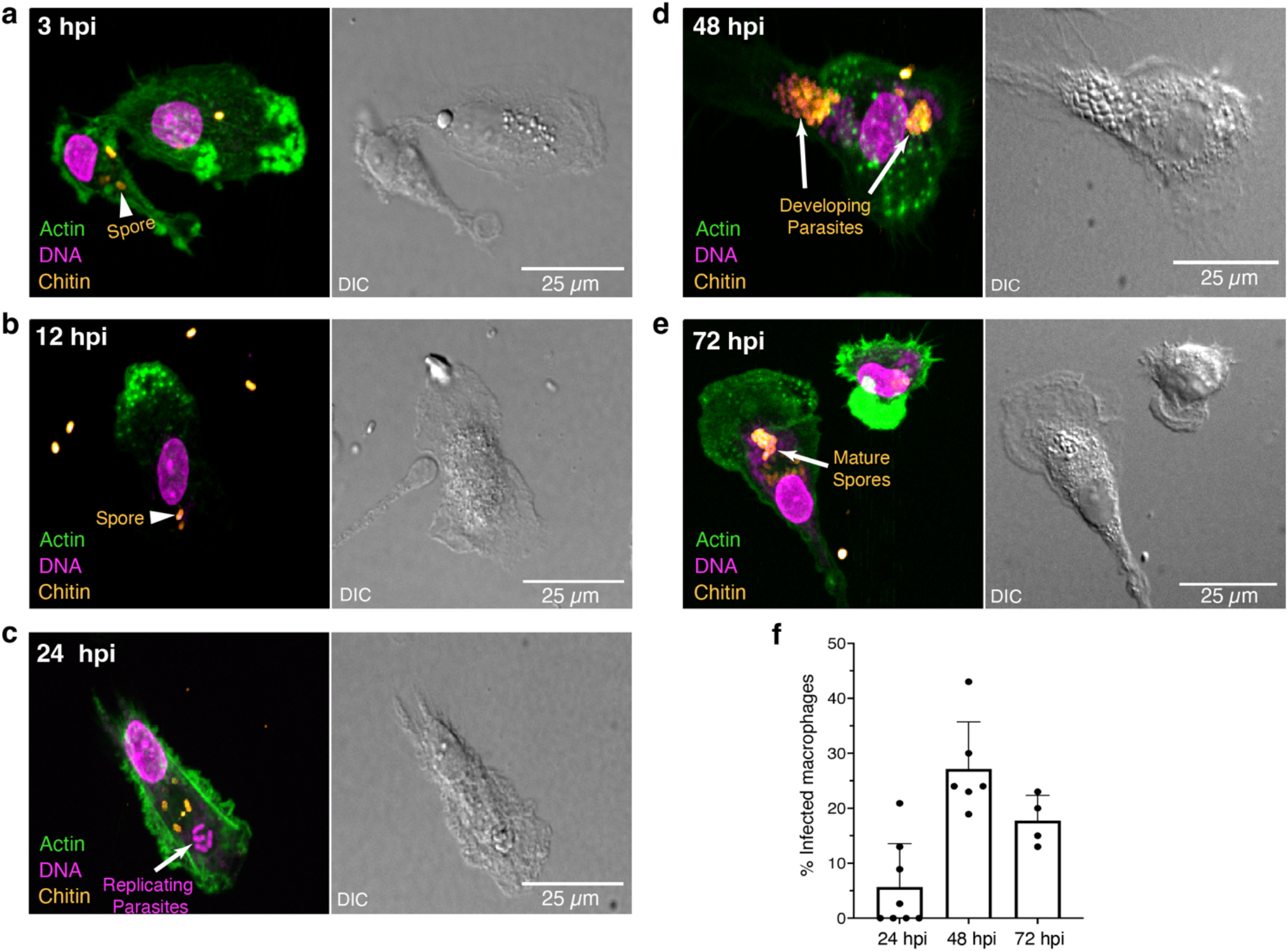
*E. intestinalis* infection kinetics in primary human macrophages. **a-e.** Representative confocal microscopy images of primary human macrophages infected with *E. intestinalis* at 3 (a), 12 (b), 24 (c), 48 (d) and 72 (e) hours post infection. A single mature spore positive for chitin (orange) depicts phagocytosis of the spore by the macrophage (arrowheads) (a-b). A cluster of DNA-only positive foci (magenta) represents parasites actively proliferating inside of the macrophage (c) (arrow). A mixture of parasites positive for DNA and chitin staining, and DNA-only staining represents macrophages with developing spores (arrow) (d-e). **f**. Quantification of macrophages with active infection. Each data point represents a biological replicate.

### scRNA-seq analysis of human macrophages infected with *E. intestinalis*

To investigate transcriptional changes in both the host and the parasite during infection, we performed scRNA-seq of *E. intestinalis*-infected macrophages. After filtering and quality control (See Methods), we analyzed a total of 39,881 cells from four healthy human donors, sampling several infection time points, alongside uninfected controls (Fig. 2a; Extended Data Fig. 1; Supplementary Table 1). We initially analyzed 8,097 macrophages from Donor 1 and 10,062 macrophages from Donor 2. Analysis of Donor 1 and Donor 2 revealed donor-to-donor variation, which prompted us to carry out further experiments on two additional donors, Donor 3 and Donor 4, with 10,254 macrophages and 11,468 macrophages, respectively.

**Figure 2.**
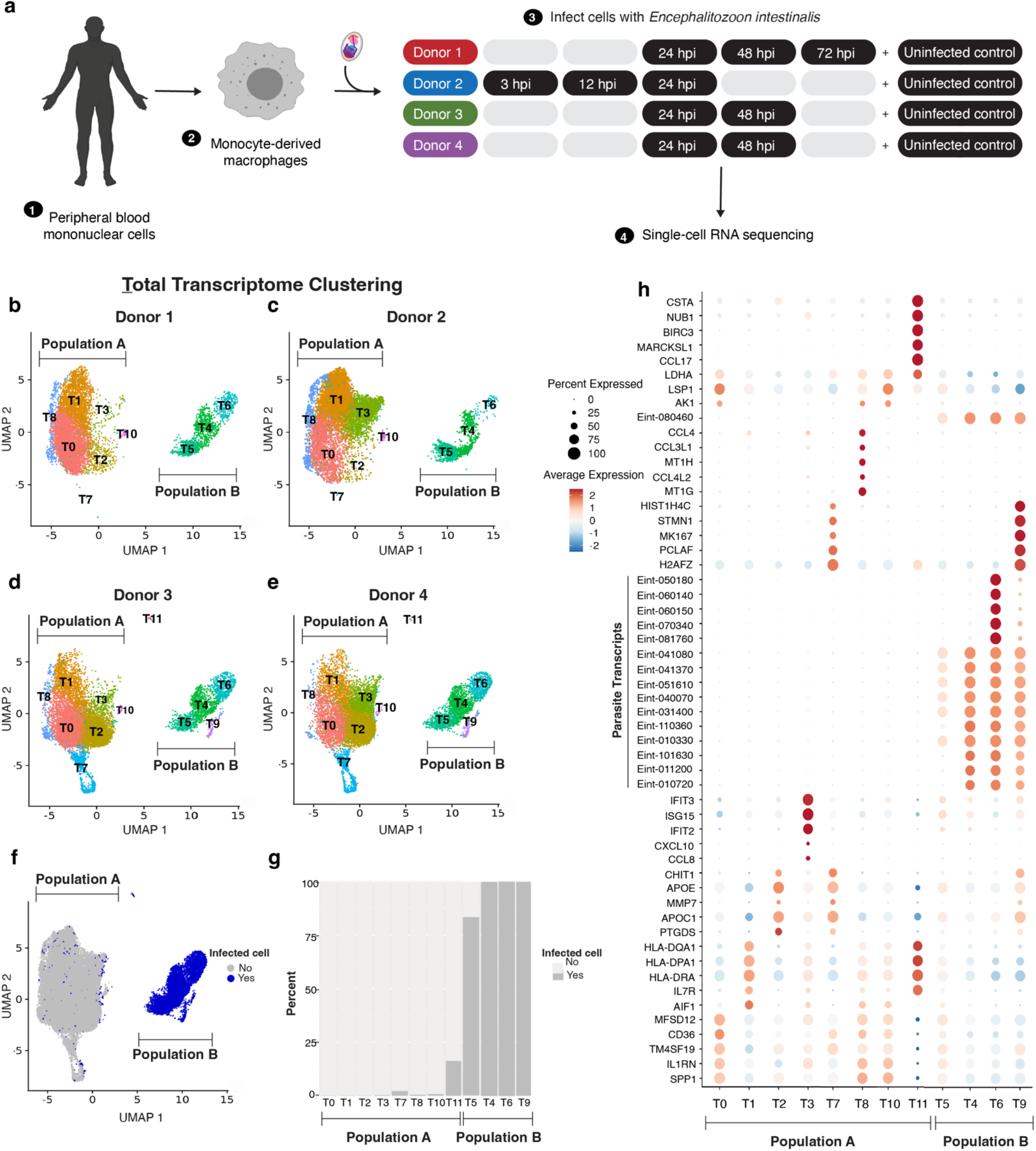
Transcriptional analysis of macrophages infected with *E. intestinalis*. **a.** Schematic overview of the experimental workflow for scRNAseq. Primary human PBMCs were collected from 4 healthy donors, differentiated into macrophages, and infected with *E. intestinalis* for 3, 12, 24, 48, or 72 hours. Uninfected cells served as controls. Cells were then prepared and taken through the scRNAseq workflow. **b-e**. UMAP plots of the integrated cells from donors projecting cells from Donors 1-4. **f**. UMAP plot of the integrated cells from Donors 1-4 colored by uninfected (gray; Population A) vs infected (blue; Population B). A cell is scored as infected if greater than 2% of total transcripts in the cell were derived from *E. intestinalis*. **g**. Quantification of the percentage of infected cells in each cluster **h**. Dotplot of the top 5 genes expressed in each cluster. The X axis shows the genes and the Y axis shows the clusters. The color indicates the average expression level across the cells in each cluster and the size indicates the percentage of cells in each cluster expressing that gene.

We first analyzed the combined parasite and macrophage transcriptomes across all four donors and all timepoints. Uniform manifold approximation and projection (UMAP) and unsupervised clustering of these cells based on highly variable genes revealed 12 clusters, T0-T11, where the “T” prefix denotes clustering based on the Total transcriptome of both the host and the parasite (Fig. 2b-e). The clusters partition into two distinct cell populations: Population A, which is present both in uninfected controls and infected samples, and Population B, which is observed almost exclusively in samples infected with *E. intestinalis* (Fig. 2b-e; Extended Data Fig. 2). To differentiate between infected and uninfected macrophages, we scored a cell as “infected” if greater than 2% of total transcripts were derived from *E. intestinalis*. We then assessed how infected cells were distributed between the two populations, and found that Population A consists primarily of uninfected cells (Fig. 2f) and includes clusters T0, T1, T2, T3, T8, and T10. In contrast, greater than 90% of the cells in Population B are infected (Fig. 2f-g) and Population B includes clusters T4, T5, T6, and T9.

Time-point analysis reveals a similar overall infection trajectory in Population B among all four donors (Extended Data Fig. 2). As early as 3 hpi, infected cells begin to populate cluster T5, and expand to also fill cluster T4 by 12 hpi. Finally, by 24 hpi, infected cells are distributed across all of the clusters in Population B (clusters T4, T5, T6, and T9), and no additional clusters arise at later time points (Extended Data Fig. 2). While samples from all donors result in a similar overall architecture and infection trajectory, there are differences in the distribution of cells between clusters. For example, Donor 2 and Donor 4 have many more cells in cluster T3 compared to Donor 1 and Donor 3 (Fig. 2b-e; Extended Data Fig. 2). Cluster T3 was largely absent from uninfected controls, but can be detected as early as 3 hpi, expands significantly by 12 and 24 hpi, then appears to decline at 48 and 72 hpi. (Extended Data Fig. 2; Supplementary Table 3). Interestingly, cells in cluster T3 are generally not infected, suggesting that these cells may be responding to the presence of *E. intestinalis* in the culture. In addition, clusters T7 and T9 are specific to Donors 3 and 4, which further suggests variation between donors, such as genetic variation or recent pathogen exposure. Alternatively, technical variation between experiments may account for some differences (see Methods). The human genes that are upregulated in clusters T7 and T9 are involved in cell cycle regulation and cell division (e.g. *H2AFZ, PCLAF, MKI67*, and *STMN1*). While clusters T7 and T9 have similar macrophage expression profiles, cells in cluster T7 are not infected by *E. intestinalis* while cells in cluster T9 are infected (Fig. 2h,g). This suggests that upon infection with *E. intestinalis*, the cells in cluster T7 give rise to the cells in cluster T9.

To investigate the gene expression signatures that define each cluster, we identified differentially expressed genes in each cluster. Clusters T4, T5, T6, and T9, which primarily contain infected macrophages, revealed that the most differentially expressed genes are from *E. intestinalis*. In clusters T4, T5, and T6 we could not detect any human transcripts with a fold change greater than 2, and in cluster T9, differentially expressed human genes are in the minority. Thus, cell clustering appears to be based primarily upon whether a cell is infected or not, while less extreme transcriptional variation in the host and parasite may be occluded and not well separated. Therefore, to gain insights into the host response to infection, as well as the developmental program in the parasite, we further analyzed the transcriptional patterns of the host and parasite separately.

### Analysis of host single-cell transcriptomes

To understand how macrophages respond to *E. intestinalis* infection, we separately analyzed only host transcripts. Cell clustering revealed 9 distinct clusters (H0-H8) (Fig. 3a; Extended Data Fig. 3a), with the “H” prefix denoting clustering based solely on the Host transcriptome. Comparison of the expression pattern for each cluster to the Human Primary Cell Atlas (HPCA)^20,21^ indicates that all of the cell clusters correspond to macrophages, monocytes, or dendritic cells (Extended Data Fig. 3b). Clusters H0, H1, H2, H4, and H7 were observed in both uninfected and infected samples, while clusters H3, H5, H6, and H8 are expanded in infected samples compared to uninfected controls. Clusters H6 and H8 contain very few cells and will not be considered further (Extended Data Fig. 3a).

**Figure 3.**
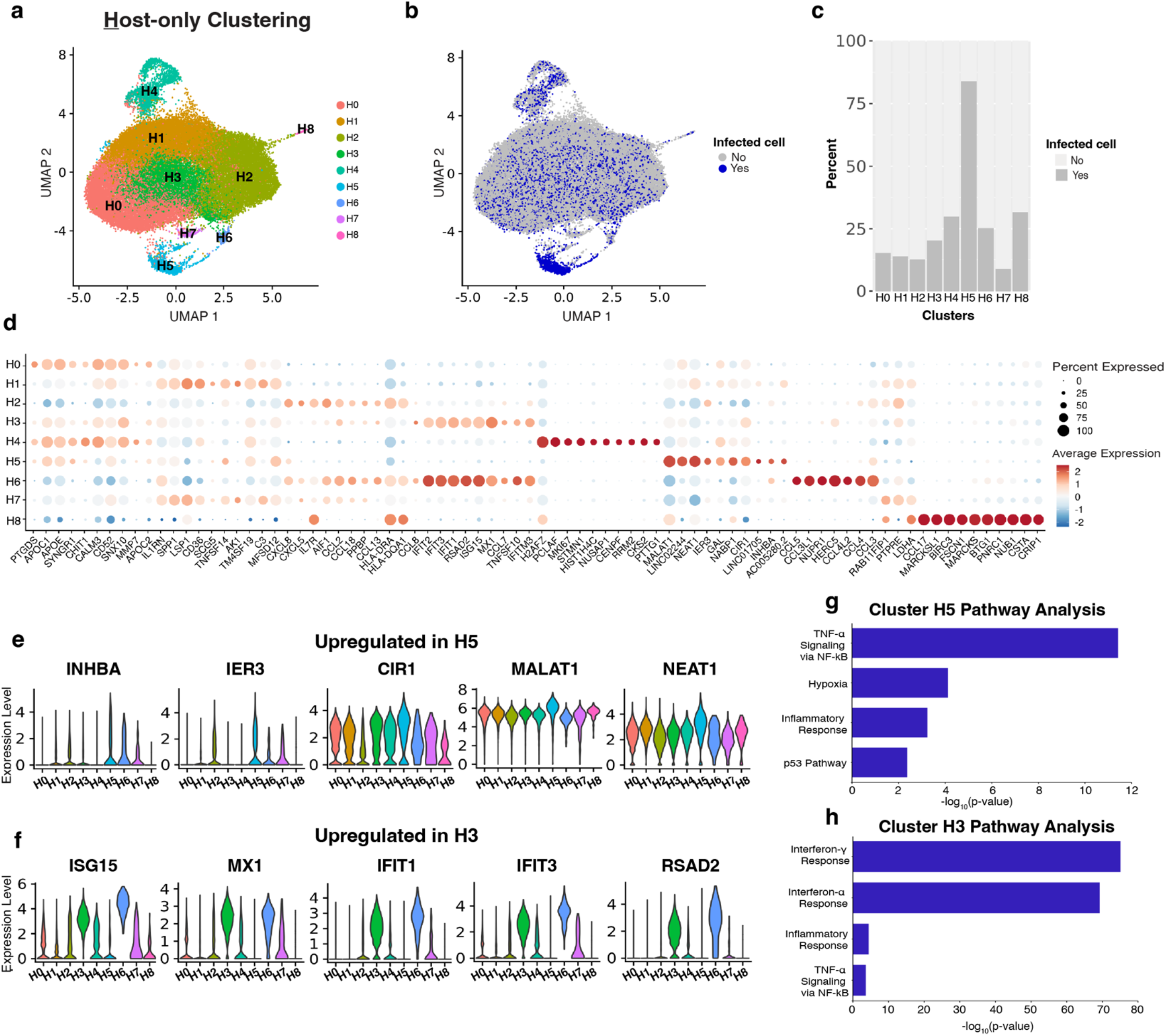
Transcriptional analysis of the host response to *E. intestinalis* infection. **a.** UMAP plot of host-only transcripts from Donors 1-4. **b**. UMAP plot colored by uninfected cell (gray) or infected (blue) cells. A cell is scored as infected if greater than 2% of total transcripts in the cell were derived from *E. intestinalis*. **c**. Quantification of the percentage of infected cells in each cluster. **d**. Dotplot of the top 10 genes expressed in each cluster. The X axis shows the genes and the Y axis shows the clusters. The color indicates the average expression level across the cells in each cluster and the size indicates the percentage of cells in each cluster expressing that gene. **e-f**. Violin plots showing expression levels of top genes from clusters H5 (e) and H3 (f). **g-h**. Bar plots showing −log_10_(p value) from enrichment analysis of biological pathways on clusters H5 and H3 using the Molecular Signatures DataBase (MSigDB) hallmark gene sets.

We next examined the distribution of infected cells across the macrophage clusters. The only cluster enriched for infected cells was H5, in which 83.9% of cells were infected. However, surprisingly, cluster H5 only contains ∼13% of all infected cells, while the majority of infected cells (87%) are distributed across the other eight clusters in approximately similar proportions (Fig. 3b), with infection rates ranging from 8.9% - 31.6%. Most infected cells are interspersed amongst non-infected cells, suggesting that in most cases, host transcription does not significantly change upon infection with *E. intestinalis*. Thus, most infected macrophages apparently fail to detect or respond to the invading parasites. In contrast, cells in cluster H5 are transcriptionally distinct from uninfected cells, and may be responding to infection. Cluster H5 appears as early as 3 hpi and persists through later timepoints. To better understand the nature of the host response in cluster H5, we identified differentially expressed genes (Fig. 3d). We observed up-regulation of cell signaling genes (such as *INHBA, IER3*), *CIR1*, and several long non-coding RNAs of known (*MALAT1* and *NEAT1*) and unknown (*LINC02244* and *LINC01705*) function (Fig. 3e). *MALAT1* and *NEAT1* have both been shown to promote inflammatory responses in macrophages during viral infection and *NEAT1* has been implicated in inflammasome activation leading to programmed cell death^22,23,24^. Pathway analysis revealed that these differentially expressed genes are involved in TNF-α signaling via NFκB, hypoxia, and inflammatory responses (Fig. 3g). The parasites found in cluster H5 are primarily at the latest stages of infection (discussed in the next section), while early parasite stages are poorly represented (Extended Data Fig. 3c). Taken together, the characteristics of macrophages in cluster H5 suggest that these cells have sensed the infection and may be responding by triggering pyroptosis to limit parasite replication and spread. Alternatively, the parasites may be manipulating the host cell in order to facilitate parasite egress via cell lysis.

Cluster H3 is also expanded in infected samples compared to controls, but in contrast to H5, H3 is not enriched for infected cells (infection rate ∼20.1%). Comparison of the H3 transcriptional profile to the Human Primary Cell Atlas revealed a strong signature characteristic of monocyte-derived macrophages that were treated with interferon-alpha (IFNα) (Extended Data Fig. 3b). Pathway analysis for genes differentially expressed in cluster H3 also indicated a strong signature related to stimulation with IFNα and IFNγ (Fig. 3H). Macrophages typically produce type I IFNs, such as IFNα, in response to viral infection, which induces apoptosis of virus-infected cells^25,26^. Therefore, it is possible that cells in cluster H3 are responding to other cells in the population that have sensed *E. intestinalis* and are secreting IFNs. In agreement with this IFNα-responsive signature, cluster H3 shows an up-regulation of interferon inducible genes such as *ISG15, MX1, RSAD2, IFIT2* and *IFIT3* (Fig. 3d,f). While the interferon response is important for the control of microsporidia infection in animal models^27^, it does not appear to be protective in macrophages *in vitro*, as parasite development appears to be unaffected in cluster H3 (Extended Data Fig. 3c).

### Analysis of *E. intestinalis* gene expression dynamics

To study the dynamics of microsporidian gene expression during parasite development, we analyzed our scRNA-seq datasets focusing only on the *E. intestinalis* transcripts. Whereas “single cells’’ in our experiment correspond to single macrophages, each infected macrophage contains one or more parasite cells, and a given cell may contain a mixture of parasite stages within one or more parasitophorous vacuoles^28,29^. Thus, the parasite mRNA within each macrophage represents the average transcriptional profile of a somewhat asynchronous population of parasites within a single macrophage. Despite these limitations, cell clustering analysis using only parasite transcripts reveals 5 distinct clusters, (Fig. 4a), potentially indicating distinct developmental stages of the *E. intestinalis* life cycle. We denote these clusters with a “P” prefix (P0-P4) since they are based solely on Parasite transcription. Two lines of evidence suggest that parasite development begins in cluster P0 and proceeds in numerical order to cluster P4. First, the percentage of microsporidian reads varies between clusters, with relatively low levels in clusters P0 and P1, and relatively high levels in clusters P2, P3, and P4 (Fig. 4b). This trend suggests that parasite burden increases between P0/P1 and P2/P3/P4 and that these correspond to earlier and later stages of the parasite life cycle, respectively. Second, time point analysis further supports this overall trajectory of infection. At 3 hpi, infected cells are found almost exclusively in cluster P0 (Fig. 4c), suggesting that this represents the initial transcriptional state immediately after host cell entry. By 12 hpi, infected cells are evenly distributed between clusters P0, P1, and P2. As cells in cluster P2 have higher levels of microsporidian mRNA compared to cluster P1 (Fig. 4b), it seems likely that parasites in cluster P2 have progressed somewhat further through the life cycle. By 24 hpi, infected cells are distributed across all five clusters (Fig. 4c), suggesting that clusters P3 and P4 represent later stages of development. Although mature spores were not observed by fluorescence microscopy until 48 hpi, the distribution of cells across clusters P0-P4 was very similar at 24, 48, and 72 hpi, with no additional populations detected at later time points. This could either be due to the resistance of hardy spores to detergent-based lysis, or due to the low RNA levels in dormant spores. Consequently, the latest stage we observe, cluster P4, may represent parasites that are not fully mature.

**Figure 4.**
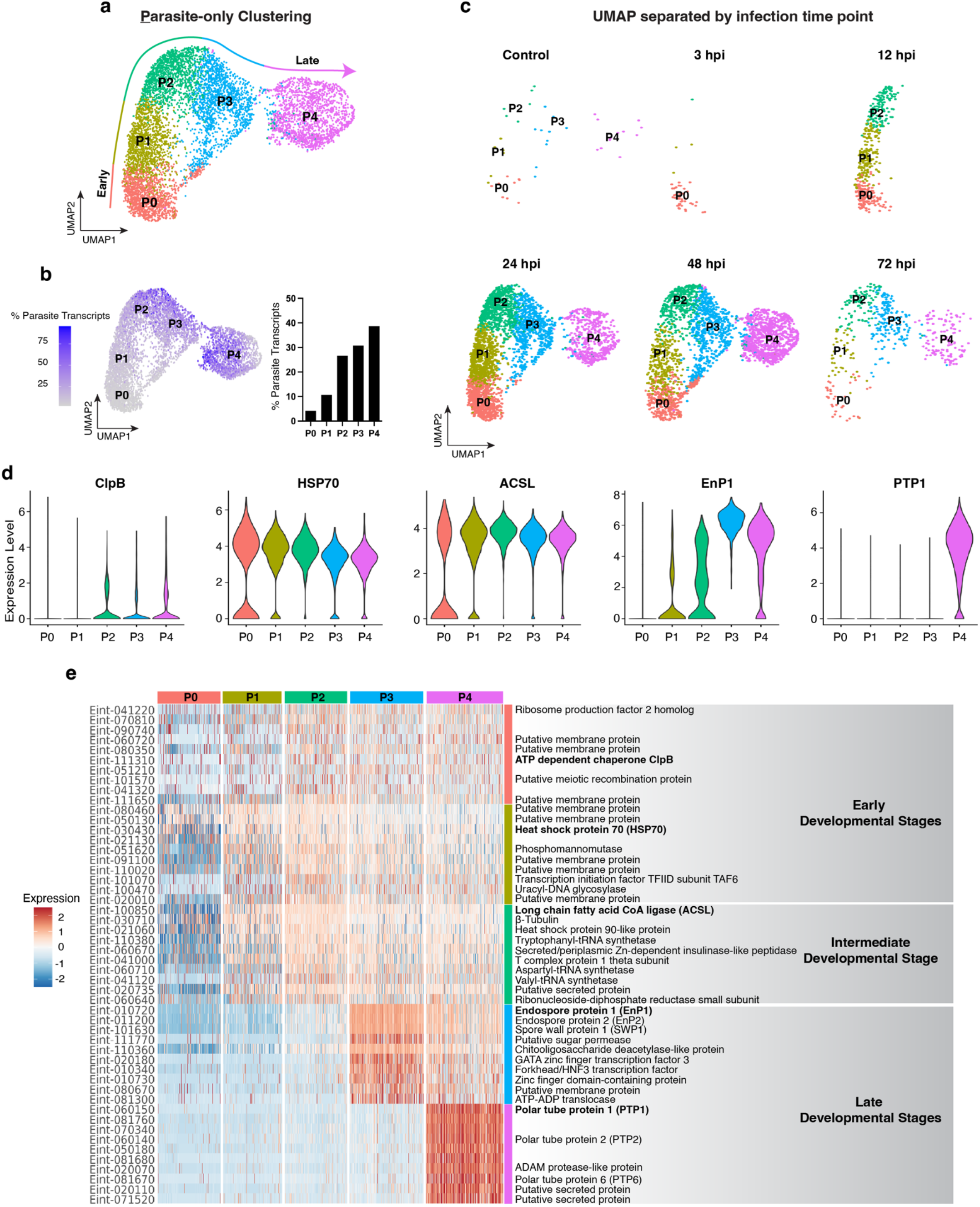
Dynamics of *E. intestinalis* gene expression during parasite development. **a.** UMAP plot of parasite-only transcripts from Donors 1-4. Each cluster corresponds to a stage of parasite development. **b**. UMAP projection of the percentage of parasite reads in a cell (left), and the percentage of parasite transcripts quantified per cluster (right) **c**. UMAP plot separated by time point reveals parasite development trajectory. **d**. Violin plots representing the expression of the top differentially expressed genes across clusters. **e**. Heatmap of differentially expressed genes in each cluster over the parasite life cycle. The color indicates relative gene expression, with red indicating increased expression. The panel on the right depicts known protein products and their role during development. The bolded protein products correspond to the genes shown in **d**.

### Differential gene expression analysis of *E. intestinalis* during development

Examination of each parasite cluster showed that very few genes were differentially expressed in clusters P0-P2, the earliest stages of infection that we detect (Fig. 4e; Supplementary Table 4). Of the genes expressed in these clusters, several are molecular chaperones and housekeeping genes such as ClpB (Eint_111310), Hsp70 (Eint_030430), and Hsp90-like protein (Eint_021060), as well as transcription initiation factors and machinery for protein synthesis (Fig. 4e; Supplementary Data 1 and 2). These housekeeping genes remain constitutively expressed throughout development, which may explain the low number of differentially expressed genes among these early developing stages (Supplementary Data 1). In cluster P2, we observed a slight increase in the expression of the long chain fatty acid coA ligase or synthetase (ACSL) (Eint_100850) (Fig. 4d,e; Supplementary Table 4; Supplementary Data 1). ACSL plays an important role in the activation of fatty acids by catalyzing the formation of fatty acyl-CoA^30^, which is required for fatty acid degradation but also as a precursor for phospholipid and triglyceride biosynthesis^31^. Hence, it is plausible that ACSL may be responsible for membrane biogenesis, which is important for the development of the endomembrane system and the rapid growth of *E. intestinalis*. Taken together, clusters P0 and P1 may represent the preparatory phases, since no other structural proteins are being expressed at this time, and P2 possibly marks the beginning of the proliferative stage.

Cluster P3 marks a shift away from fundamental cellular metabolism and proliferation, towards more specialized functions required for the production of spores (Fig. 4e). The magnitude of transcriptional changes is up to 6-fold in P3, compared to the more modest changes in clusters P0, P1, and P2, which reach a maximum of 2-fold (Fig. 4e; Supplementary Table 4; Supplementary Data 2). The two most upregulated genes in cluster P3 are Endospore Proteins 1 and 2^32^ (EnP1; Eint_010720 and EnP2; Eint_011200, respectively), along with several other proteins thought to be localized to the spore coat, such as Spore Wall Protein 1 (SWP1; Eint_101630), as well as proteins involved in the synthesis and remodeling of cell wall polysaccharides, such as a putative Chitooligosaccharide Deacetylase (Eint_110360) and Chitin Synthase (Eint_011330) (Fig. 4e; Extended Data Fig. 4; Supplementary Data 2). Several septins are also upregulated in cluster P3 (Eint_111850, Eint_011310, and Eint_090780). Septins are often involved in creating boundaries between cellular compartments, such as between the cell body and the cilium^33^ or between the parent cell and the daughter bud in *Saccharomyces cerevisiae*^*33,34*^, but their functions in microsporidia remain poorly characterized. Several transporter and aquaporin genes are also upregulated (Eint_111770, Eint_081300, and Eint_070680), which may be important for siphoning key nutrients from the host and osmoregulation during parasite maturation (Fig. 4e; Extended Data Fig. 4; Supplementary Table 4; Supplementary Data 2). Finally, several transcription factors are upregulated, which may be involved in the transition from P2 to P3 or from P3 to P4.

At the latest stage of development, cluster P4, we observed the largest changes in gene expression, up to 41-fold, including upregulation of many known components of the polar tube, a harpoon-like invasion apparatus assembled in maturing microsporidian spores (Fig. 4e; Extended Data Fig. 4; Supplementary Table 4). Polar Tube Protein 1 (PTP1; Eint_060150), is the most highly upregulated gene in cluster P4 (41-fold), and is the most differentially expressed gene across all 5 parasite clusters (Supplementary Table 4; Supplementary Data 2). Also upregulated in cluster P4 are the known Encephalitozoon polar tube proteins PTP2 (Eint_060140), PTP3 (Eint_111330), and PTP4 (Eint_071050); and the proposed ortholog of *N. bombysis* PTP6 (Eint_081680) (Extended Data Fig. 4; Supplementary Data 2). Thus, cluster P4 appears to include several genes known to encode components of the polar tube, and may also include additional genes important for polar tube assembly and spore development, which occurs during the mid-to-late-sporoblast stage^29^.

### Clusters P3 and P4 are enriched in secreted proteins involved in spore maturation

The structures of most PTPs are poorly predicted by AlphaFold2 and have minimal similarity to known protein domains. However, PTP4 and PTP6 both contain a single Ricin B-type Lectin domain (RBL)^35,36^, and other RBL proteins have also been linked to the PT^37^. Previous work has suggested that RBL domains form a small, beta-trefoil domain involved in carbohydrate binding^38^, and that PTP4 in particular may play a role in attachment to host cells during invasion^35^. Hypothesizing that other RBL proteins may also play a role in PT function, and therefore be transcriptionally co-regulated, we searched the *E. intestinalis* genome for genes encoding RBL domains and assessed if they were also upregulated in cluster 4. We identified a total of 14 putative RBL proteins in *E. intestinalis* (see Methods), including PTP4, PTP6, and a protein that has been referred to as “PTP5” in unpublished reports and reviews^39^ (Extended Data Fig. 5). Of these 14 putative RBL proteins, 12 are potential PTP4/PTP6 paralogs, and appear to encode a single RBL domain with a potential N-terminal signal peptide to direct them to the secretory pathway (Fig. 5a); the remaining 2 putative RBL proteins are architecturally distinct, and appear to consist of an RBL domain fused to an integral membrane protein and are annotated as putative Dolichyl-phosphate-mannose protein O-mannosyltransferases (Fig. 5a). 12 of the 14 putative RBL genes, including 10 of the 12 PTP4/PTP6 paralogs, are upregulated in cluster P4 (Fig. 5b). The remaining 2 PTP4/PTP6 paralogs were not upregulated in any cluster, and expression levels are low. Taken together, there is a clear and coordinated upregulation of PTP4/PTP6-like RBL proteins in cluster P4, which may indicate a more general role for this protein family in polar tube assembly and/or spore maturation. In addition to PTPs and RBL proteins, cluster P4 shares some of the signatures upregulated in cluster P3, including septins, and several genes involved in the synthesis of cell surface polysaccharides.

**Figure 5.**
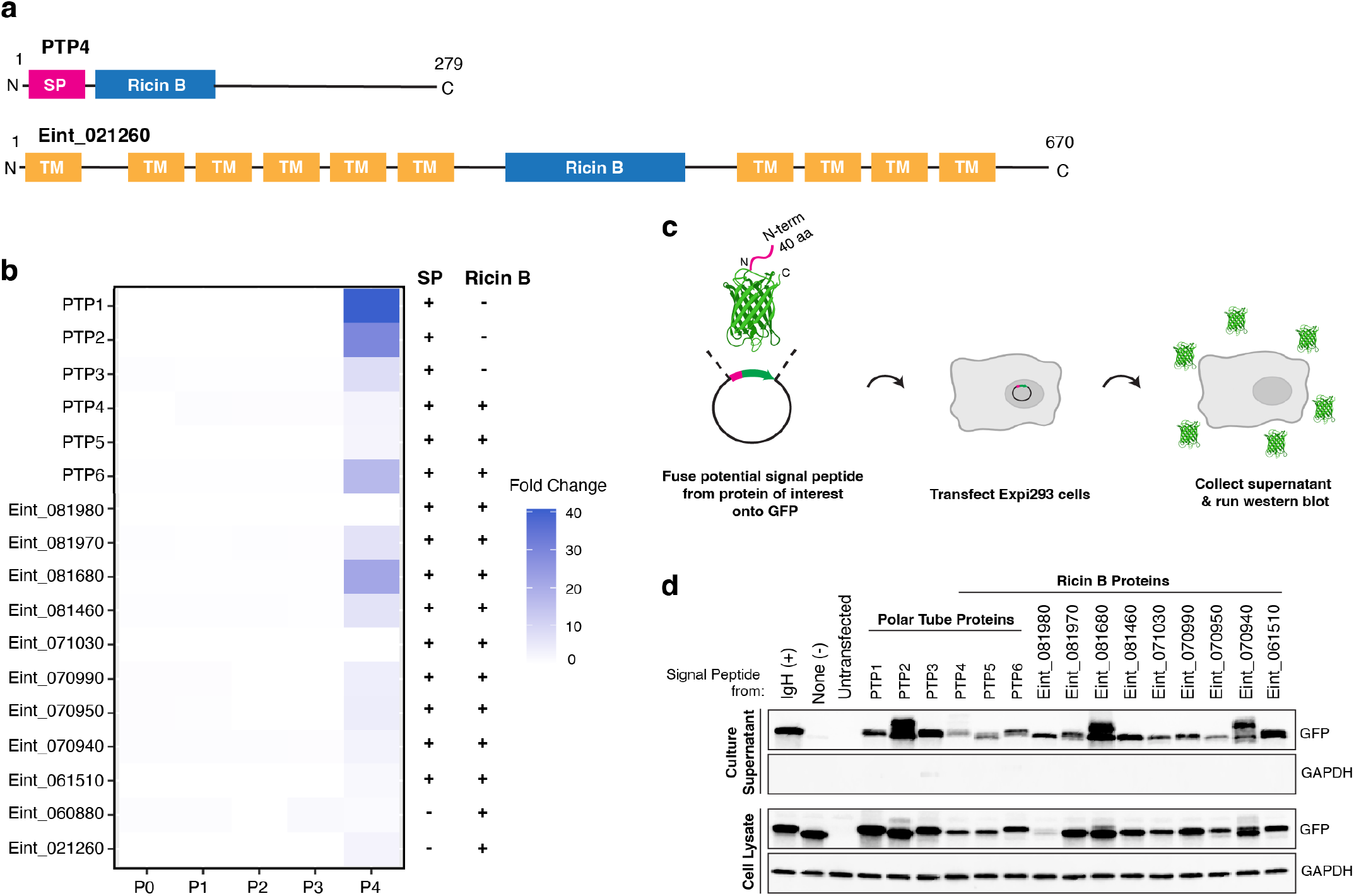
Cluster P4 is enriched in proteins containing signal peptides and Ricin-B domains. **a.** Schematics of representative proteins containing a Ricin-B domain and an N-terminal signal peptide (top) and proteins containing a Ricin-B domain fused to an integral membrane protein (bottom). **b**. Heatmap depicting the expression of genes in cluster P4 containing a Ricin-B domain and/or a signal peptide. **c**. Schematic of the assay we developed to test secretion. Constructs containing the first 40 amino acids of each protein from **b** fused to the N-terminus of GFP were transfected into Expi293 cells, alongside control constructs. Supernatants and whole cell lysates were collected and analyzed via western blot, including lysis controls **d**. Western blot of supernatants (top) and whole cell controls (bottom) from panel C, probing GFP and GAPDH.

Many of the known proteins upregulated in clusters P3 and P4 are targeted to the secretory pathway, including the PTPs^40^ and SWP1^41^. Numerous other proteins of unknown function are also upregulated in clusters P3 and P4, leading us to hypothesize that some of these might also play roles in cell wall or PT assembly/function and, therefore, might also be targeted to the secretory pathway. Thus, we assessed whether genes upregulated in clusters P3 and P4 were enriched for secreted proteins relative to earlier developmental stages, and to the *E. intestinalis* proteome as a whole. We used SignalP-6.0 with a relaxed cutoff to compensate for the insensitive detection of signal peptides in highly divergent microsporidian proteins (see Methods). Using this approach, we predict that ∼5.6% of *E. intestinalis* proteins may be secreted (108 of 1934 proteins) (Supplementary Table 5, Extended Data Fig. 6). Relative to the *E. intestinalis* proteome at large, clusters P0, P1, and P2 are not enriched for secreted proteins (1.7%, 3.7%, and 6.8%, respectively). The frequency of proteins with predicted signal peptides jumps to 9.6% in cluster P3, and 24% in cluster P4. The upregulated genes in cluster P4 encode nearly half of all the predicted secreted proteins in the *E. intestinalis* proteome. Of the 47 putative secreted proteins upregulated in cluster P4, 15 are PTPs or RBL proteins (Fig. 5b). To test whether these 15 proteins are secreted by *E. intestinalis*, we implemented a protein secretion assay in a heterologous, mammalian expression system. We designed constructs in which the N-terminal 40 residues of each protein, including the putative signal peptide, was fused to the N-terminus of GFP. We then transfected each construct into Expi293 cells, and assessed the secretion of our GFP fusion proteins into the culture supernatants 24 hrs after transfection (Fig. 5c). Western blotting against GFP revealed that the N-terminus of all 15 proteins is capable of mediating GFP secretion from mammalian cells (Fig. 5d), suggesting that they likely encode bonafide signal peptides that target the native proteins to the secretory pathway in *E. intestinalis*. In addition to these 15 PTP and RBL proteins, 4 of the 47 proteins that are upregulated in P4 and predicted to be secreted are proteases, which may be involved in processing and maturation of secreted proteins. Most of the remaining 28 secreted proteins have poorly predicted structures using AlphaFold2 and lack clear similarity to proteins of known structure or function. However, the strong upregulation of many of these secreted proteins, along with known PTPs, suggests possible roles in polar tube biogenesis or spore maturation.

**Figure 6.**
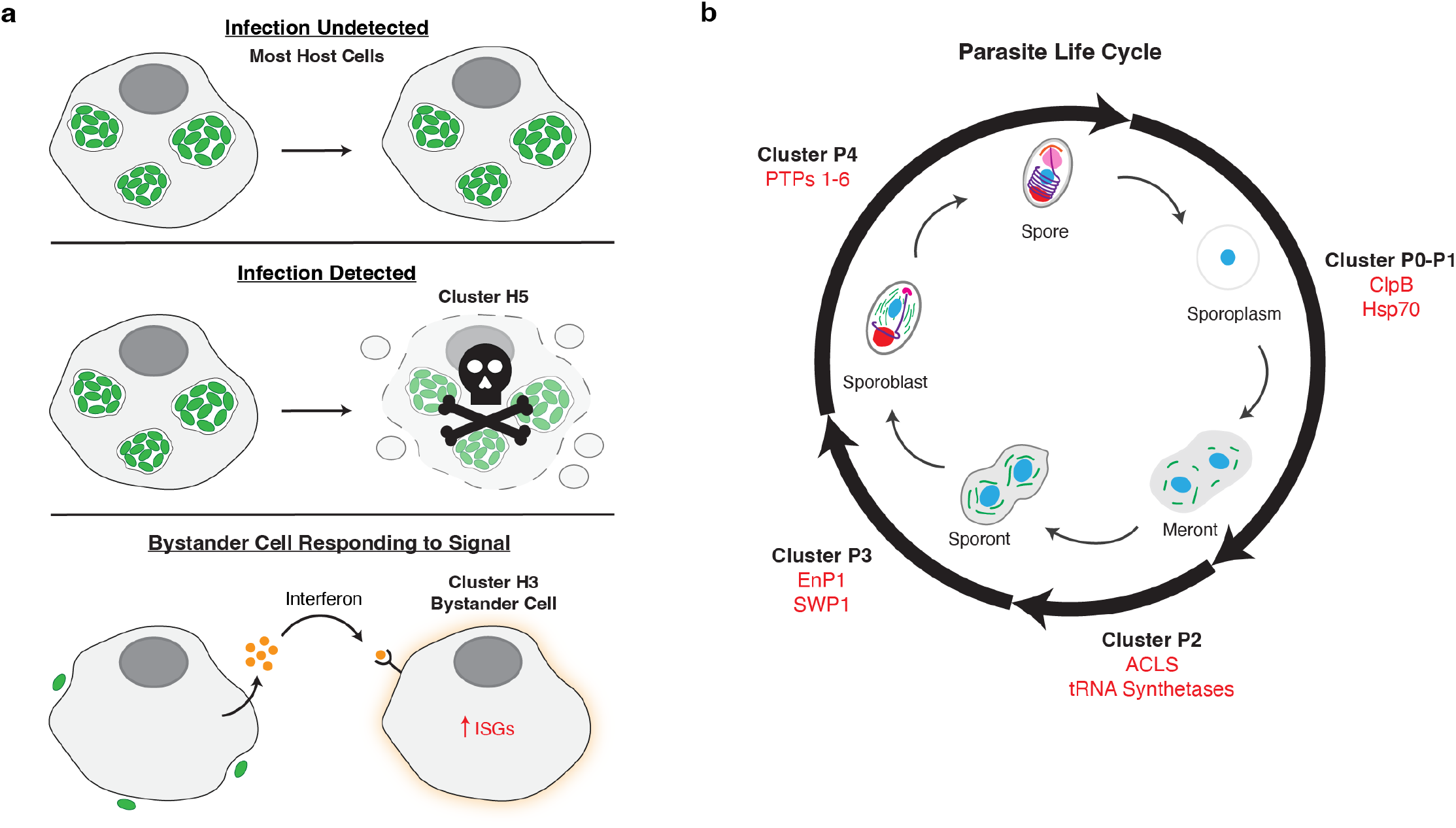
Host-parasite transcriptional atlas. **a.** Schematics of the host response to *E. intestinalis* infection. *E. intestinalis* infection goes undetected in the majority of cells and the parasites are able to undergo their full lifecycle (top). A subset of host cells are able to respond to infection (cluster H5) particularly when the parasites mature into late stage spores (middle). These cells are undergoing cell death to limit infection and spread. Bystander cells are responding to secreted interferons from infected cells and creating an antiviral cell state to try to prevent infection (bottom). **b**. Developmental program of *E*. intestinalis. Molecular markers for each parasite stage in red.

## DISCUSSION

The transcriptional response to microsporidia infection among hosts are diverse. Previous studies using bulk RNA seq found in silkworms infected with *Nosema bombycis* there is an upregulation of antimicrobial peptides^42^ and *Caenorhabditis elegans* undergo an antiviral response when infected with *Nematocida* species^42,43^. In mammals, cytokines are induced upon infection and there is an overall rewiring of host cell energy production^18,44^. Here, we investigated single cell transcriptomic changes during *E. intestinalis* infection in primary human macrophages from four healthy human donors. Our results indicate that the majority of macrophages are unable to detect invading *E. intestinalis* parasites, as depicted by the even distribution of both infected and uninfected macrophages across clusters (Fig. 3b; Fig. 6a). In most macrophage clusters, we observed all parasite developmental stages, which suggests that *E. intestinalis* can progress through its entire life cycle without altering the transcriptional profile in host macrophages and successfully avoid host detection. This is in agreement with previous studies, in which naive human monocyte-derived macrophages were unable to reduce parasite burden^45^. However, if monocyte-derived macrophages were pretreated with LPS or interferon-γ, they were able to reduce parasite burden, suggesting that naive macrophages require external signals, often from T cells, in order to inhibit parasite replication. This mimics the environment in immunocompromised individuals, such as AIDS patients, where there is a decreased number of T cells to provide external signals to the infected macrophages, allowing *E. intestinalis* to harness macrophages for replication and dissemination. While the majority of cells are unable to detect infection, one cluster, H5, is induced upon infection. Cluster H5 contains a high rate of infected cells and is transcriptionally distinct from other clusters (Fig. 3b), suggesting that cells in this cluster detect infection and may mount an inflammatory response against the parasites. Interestingly, the parasites in this cluster primarily represent the latest stages of development (Extended Data Fig. 3c). It is possible that mature parasites are easier to detect, resulting in the host cell undergoing pyroptosis to limit infection (Fig. 6a). Like H5, cluster H3 is also induced by *E. intestinalis*. However, in contrast to H5, the rate of infection in H3 is much lower, and cells from H3 may be responding to interferons that are being secreted by neighboring infected cells instead of responding directly to infection (Fig. 6a).

An advantage of scRNA-seq is that it simultaneously captures both host and parasite transcriptomes, allowing us to monitor the developmental program of *E. intestinalis*. Our data reveal a transcriptional atlas of *E. intestinalis* development, which enables the identification of molecular markers for each stage of parasite development. During the early stages of *E. intestinalis* development (P0-P1), we see house-keeping genes being expressed as well as transcription initiation factors. This suggests that clusters P0-P1 represent parasites just prior to replication and rapid proliferation, which may correspond to the sporoplasm or meront stages (Fig. 6b). In cluster P2, we observe the expression of ASCL as well as protein synthesis machinery which are likely important for the rapid growth of *E. intestinalis*, corresponding to the proliferative meront to early sporont stages. Following the proliferative stage, chitooligosaccharide deacetylase-like protein was up-regulated (Fig. 4e). This protein is involved in deacetylation of chitin to form chitosan, an important component of the fungal cell wall^46^. It has been shown in *E. cuniculi* that the chitin deacetylase localizes to the plasma membrane during the meront-to-sporont transition^47^. This suggests that cluster P3 begins the development of the spore wall. This was further exemplified by the up-regulation of genes involved in spore wall formation including SWP1 and EnP1. In *E. intestinalis*, SWP1 is expressed in the transition between the proliferative stage and sporont stage, in which a thin layer of exospore is formed, suggesting that P3 is the sporont stage. In the final detected stage of development, the genes encoding the known PTPs are differentially expressed along with genes encoding proteins with RBL domains and signal peptides. The upregulation of many of these secreted proteins along with known PTPs suggests this cluster corresponds to PT biogenesis. Thus, cluster P4 may correspond to the mid-to-late sporoblast stage which is where PT assembly occurs (Fig. 6b). Overall our work begins to uncover the molecular mechanisms of parasite development that correspond to the developmental stages observed by previous EM studies^29^, as well as the heterogeneity of host cell responses and donor-to-donor variation.

## METHODS

### Isolation of primary human monocyte-derived macrophages

Primary human peripheral blood mononuclear cells (PBMCs) from anonymous, healthy donors (New York Blood Center) were isolated by Ficoll gradient separation as previously described^48^. CD14^+^ monocytes were then isolated from the PBMC fraction using the EasySep™ Human CD14 Positive Selection Kit II (STEMCELL Technologies) according to the manufacturer’s protocol. Briefly, the cells were resuspended at a concentration of 1x10^8^ cells/mL in PBS (Gibco) containing 2% heat-inactivated fetal bovine serum (FBS) and 1 mM Ethylenediaminetetraacetic acid (EDTA). The EasySep Human CD14 Positive Selection Cocktail II was added (100 μL/mL of sample) and incubated for 10 minutes at room temperature. Next, the EasySep™ Dextran RapidSpheres was added (100 μL/mL of sample) and incubated for 3 minutes at room temperature. The mixture was topped to the recommended volume with PBS containing 2% FBS and 1 mM EDTA. The sample was placed in an EasySep magnet at room temperature for three minutes. The supernatant was discarded and repeated for a total of three washes. The isolated CD14^+^ cells were plated at 1×10^6^/mL in RPMI (Cytivia) supplemented with 10% FBS, 10 mM HEPES, 100 U/mL penicillin, 100 μg/mL streptomycin and 50 ng/mL granulocyte-macrophage colony-stimulating factor (GM-CSF) (Partner Therapeutics). The cells were incubated at 37°C with 5% CO_2_. Media was replenished with GM-CSF on day 2. Cells were harvested and plated for infection on day 4. Donors 1 and 2 were isolated on two separate days whereas Donors 3 and 4 were isolated in parallel on the same day. Consequently, the UMAP clustering similarities among Donors 3 and 4 and the UMAP clustering differences between Donors 3 and 4 compared to Donors 1 and 2 may reflect biological difference between donors, or could also result from batch effects during cell isolation and culture, which may lead cells processed on different days being transcriptionally more similar to cells isolated on the same day.

### Propagation and purification of *E. intestinalis*

All mammalian and parasite lines used in this work were tested monthly for mycoplasma contamination (MycoStrip, InvivoGen), and all tests were negative. *E. intestinalis* spores (ATCC 50506) were propagated in Vero cells (ATCC CCL-81). Vero cells were grown in a 75 cm^2^ tissue culture flask using Dulbecco’s Minimum Essential Medium (DMEM) (Gibco) supplemented with 10% heat-inactivated fetal bovine serum (FBS) at 37°C and with 5% CO_2_. At 70%-80% confluence, the media was switched to DMEM supplemented with 3% FBS and *E. intestinalis* spores were added. Infected cells were allowed to grow for 8-12 days and the medium was changed every two days. To purify spores, the infected cells were detached from tissue culture flasks using a cell scraper and moved to a 15 mL conical tube, followed by centrifugation at 1,300 g for 10 min at 25°C. Cells were resuspended in 5 mL 1X PBS and mechanically disrupted using a G-27 needle. The released spores were purified using a 2-step Percoll gradient. Equal volumes (5 mL) of spore suspension and 100% Percoll were added to a 15 mL conical tube, vortexed, and then centrifuged at 1,800 g for 30 min at 25°C. The spore pellets were washed three times with 1X PBS. The spores were further purified via Percoll gradient (25%, 50%, 75%, 100%) ultracentrifugation. The spores were layered over the Percoll gradient and centrifuged at 8600 rpm for 30 minutes at 25°C (TH-641 Swinging Bucket Rotor, ThermoFisher Scientific). Bands containing various impurities remained in the supernatant/gradient fractions, and were removed using a vacuum. The pellet at the bottom of the tube, containing mature spores, was washed with 1X PBS and ultracentrifuged again at 8600 rpm for 30 minutes at 25°C. The spore pellet was resuspended in 1X PBS and transferred to a 1.5 mL microcentrifuge tube and centrifuged at 3,000 g for 3 minutes at 25°C. This was repeated twice. The final pellet was resuspended in 1X PBS and stored at 4°C.

### *E. intestinalis* infection experiments

For each infection experiment used to prepare the samples for scRNA-seq, two identical sets of cultures of *E. intestinalis*-infected Vero cells were prepared. One set of cultures was processed for scRNA-seq and the second set of cultures was prepared for fluorescence microscopy to assess the infection levels of the sample. 2x10^5^ differentiated macrophages were seeded in a 6-well plate (CELLTREAT), either empty (for scRNA-seq) or with 5 rounded-glass coverslips placed in each well (for microscopy sample) (Fisher Scientific). The wells were supplemented with RPMI 1640 medium, 10% FBS, and 50 ng/mL of GM-CSF. The cells were allowed to rest for 24 hr at 37°C with 5% CO_2_. 2.5 hr prior to infection with *E. intestinalis*, GM-CSF was added to the cells. To infect the cells, purified, mature *E. intestinalis* spores were added into each well at a multiplicity of infection (MOI) of 20 (or 4x10^6^ spores). Microsporidian spores were allowed to infect the cells for 24 hr, except 3hpi and 12 hpi timepoints which were incubated for 3 hr and 12 hr, respectively. For samples intended for analysis 24, 48, and 72 hpi, spores were removed and the wells were washed with fresh media 3 times at 24 hpi and infection continued to the respective timepoints. The exact infection timepoints collected in each donor is listed in Fig. 2a and Supplementary Table 1.

Fluorescence microscopy was used to assess microsporidian infection and replication in macrophages, prior to scRNA-seq. The media was removed from the wells and the cells were fixed with 4% paraformaldehyde at 37°C for 15 min, followed by washing with 1X PBS, three times. The cells were stained with 1X phalloidin-iFluor488 (Abcam, USA) at RT for 20 min. After washing with 1X PBS, the cells were stained with 5 μM DRAQ5 (Novus Biologicals) for 10 min and 0.5 μg/ml of calcofluor white for 10 min (Sigma Aldrich) at RT. To mount the slides, 50% glycerol was added to poly-lysine coated slides (Fisher Scientific) and the coverslips were placed on the slide. The coverslips were sealed with a clear nail polish to prevent evaporation. The slides were visualized using a Nikon CSU-W1 spinning disk laser confocal microscope.

### Generating single-cell suspensions and cell hashing

To isolate a single cell for scRNA-seq, infected macrophages were detached from the 6-well plate by trypsinization. Briefly, the media was removed from the well and incubated with 0.25% trypsin (Corning) at 37°C with 5% CO_2_ for 15 min. RPMI 1640 medium supplemented with 10% FBS was added to stop the trypsinization reaction. At this step, some of the macrophages still adhered to the well. Pipetting of the media against the bottom of the well was done to further dissociate the cells. To collect the cells, they were centrifuged at 1,200 g for 5 min at 4°C.

To combine cells from several infection timepoints for the scRNA-seq library preparation, the cells from each infection timepoint were incubated with different hashing antibodies^49^. Macrophages were resuspended with 100 μl of a staining buffer (2% BSA and 0.01% Tween 20 in 1X PBS), supplemented with 10 μl of Fc blocking reagent (BioLegend, USA), and incubated for 10 min at 4°C. 1 μl of TotalSeq™ B anti-human hashtag antibodies 5-10 (BioLegend, USA) was added into the cell suspensions collected from each infection time point and incubated for 20 min at 4°C. The cells were washed three times with 1 ml of staining buffer. Finally, the cell pellets were resuspended with 150 μl of staining buffer. Cell density and viability were counted using a hemocytometer by mixing the cells with trypan blue dye (Invitrogen, USA). Our cell viability in each infection time point ranges from 91% to 100%.

### Library preparation and 10X scRNA-seq

After cell hashing, cells from different infection timepoints were pooled together and adjusted to ∼500 cells/μl prior to processing for scRNA-seq library preparation. The library preparation was carried out using a Chromium Single-Cell 3′ Reagent Kits v2 Chemistry (10x Genomics, USA) according to the manufacturer’s protocol. We recovered a total of 10,100 cells from Donor 1, 12,279 cells from Donor 2, and 30,355 cells combined for Donors 3 and 4 for scRNA-seq library preparation. After library completion, the paired-end sequencing was performed using an Illumina NovaSeq 6000 system (Illumina, USA). Both library preparation and sequencing were carried out at the Genome Technology Center, NYU Langone Health.

### Processing of the raw sequencing reads

A combined reference genome containing both the human genome (GRch38)^50^ and the *E. intestinalis* genome (ATCC 50506)^51,52^ was generated using Cell Ranger software version 5.0.1 with a Cell Ranger *mkref* function (10X Genomics, USA). Raw sequencing reads were mapped to the combined reference genome and the gene expression matrices were generated using Cell Ranger *count* function with default parameters^53^. Cell Ranger *count* quantifies an amount of genes detected in each cell (nFeature) and the total number of the RNA molecules in each cell (nCount), which was evaluated from unique molecular identifiers (UMIs).

### scRNA-seq data processing

#### Initial data processing

The gene expression matrices from Cell Ranger *count* were transferred and processed using Seurat version 4.0^54^. The number of cells obtained from each donor prior to filtering are shown in Supplementary Table 6. Cells from different infection timepoints were demultiplexed based on their hashing antibodies, using a MULTIseqDemux function. Doublet cells, which contained more than one type of hashing antibodies, and negative cells, which lacked any hashing antibody) were removed from all of the datasets. To further filter low quality cells, nFeature, nCount, and percentage of mitochondrial genes (percent.mt) were used (Supplementary Fig. 2). Criteria for filtering cells were different between donors. In donor 1 dataset, the criteria were nFeature: 500-6,000 genes, nCount: 2,000-40,000 UMIs, and percent.mt < 20. For donor 2, the criteria were nFeature: 500-6,000 genes, nCount: 2,200-35,000 UMIs, and percent.mt < 20. For donor 3, the criteria were nFeature: 400-4,000 genes, nCount: 1,000-20,000 UMIs, and percent.mt < 20. For donor 4, the criteria were nFeature: 400-3,800 genes, nCount: 1,000-15,000 UMIs, and percent.mt < 20. Number of cells passed the quality control are shown in Supplementary Table 1.

The datasets for donor 1, donor 2, and donor 3+4 combined were each normalized using a log normalization method (NormalizeData, normalization.method = “LogNormalize”, scale.factor = 10,000) and variable features were identified (FindVariableFeatures, selection.method = “vst”, nfeatures = 3,000). Integration anchors were identified prior to combining the data from different donors (FindIntegrationAnchors). These anchors were used to integrate datasets from 4 donors (IntegrateData) and the integrated data were scaled (ScaleData). Then, we ran RunPCA. PC1 to PC41 were included to perform dimensionality reduction using a Uniform Manifold Approximation and Projection (UMAP) method (RunUMAP, dims = 1:41). For cell clustering, FindNeighbors (dims = 1:41) and FindClusters (resolution = 0.4) were carried out. DimPlot function was used to generate UMAP plots in Fig. 2b-f, while ggplot was used to make Fig. 2g.

To identify which macrophages were infected with *E. intestinalis*, the percentage of *E. intestinalis* transcripts detected in each cell were calculated (PercentageFeatureSet, pattern = “^Eint-”). We classified macrophages that have more than 2% of the microsporidian transcripts as “infected”. The number of infected cells found in each donor are shown in Supplementary Table 2. A small portion of the control cells are classified as infected. We suspect that, after pooling the uninfected control cells with infected samples following cell hashing, spores from infected samples may have invaded or adhered to control cells, resulting in a small number of control cells containing *E. intestinalis* transcripts.

#### scRNA-seq data processing of the *E*. *intestinalis* transcripts

To investigate *E. intestinalis* transcriptional profiles at different developmental stages, *E. intestinalis* transcripts were separated from the human transcripts by a subset function prior to normalization using identical parameters as previously described in the initial scRNA-seq data processing. Only infected cells identified from the initial scRNA-seq data processing were used in this analysis. Integration anchors were identified (FindIntegrationAnchors). The data from 4 donors were integrated (IntegrateData) and scaled (ScaleData). RunPCA was performed followed by RunUMAP using PC1 to PC44 (RunUMAP, dims = 1:44). To cluster the infected cells, FindNeighbors (dims = 1:44) and FindClusters (resolution = 0.35) were performed. FeaturePlot function was used to generate Fig. 3b,e.

#### scRNA-seq data processing of the human transcripts

To process the human transcripts separately, low quality cells were filtered using the same parameters as described in the initial data processing section. Then, the data from 4 donors were integrated and scaled. For the downstream analyses, similar pipelines as in the processing *E. intestinalis* transcripts were utilized, except the principle components used for RunUMAP. They were PC1 to PC43 (RunUMAP, dims = 1:43) and the resolution used for performing cell clustering was 0.20 (FindClusters, resolution = 0.20).

#### Marker gene detection and differential gene expression analyses

To identify signature genes for each cell cluster, FindAllMarkers function was performed using the default parameters and the min.cpt of 0.25 (FindAllMarkers, min.pct = 0.25). Wilcoxon Rank Sum test was used to test the differentially expressed genes between clusters. Average Log_2_ Fold change (avg_log2FC) was also used to rank highly expressed genes in each cluster. DotPlot was carried out to generate Fig. 2h and Fig. 3d, while DoHeatmap was utilized in Fig. 4e. VlnPlot was used to visualize the gene expression levels in Fig. 3e,f and 4d.

#### Gene ontology enrichment analysis

To gain better understanding on what pathways highly expressed genes in cluster H5 and H3 are involved in, pathway enrichment analysis was performed using EnrichR (https://maayanlab.cloud/Enrichr/)^55^. Briefly, highly expressed genes that are up-regulated and have a fold change more than 1.5 were selected and subjected for EnrichR. Fig. 3g and h were obtained using a ‘Molecular Signatures Hallmark 2020’ library. The terms were ranked according to their p-values.

#### Automatic cell type annotation

To perform automatic cell type recognition from the scRNA-seq data, SingleR package was used^21^. SingleR compares gene expression patterns from each cell cluster to the normalized expression values obtained from the Human Primary Cell Atlas (HPCA)^20^. A ‘label.fine’ function was used to obtain more detailed categories of the immune cell types (SingleR, ref = Human.primary.cells, assay.type.test = 1, labels = Human.primary.cells$label.fine). Each cell in the datasets was categorized into one of the cell types. Then, the percentage of cells in each cluster were visualized using ggplot in R.

### Prediction of Ricin B domain-containing proteins

To identify proteins with ricin B domains in *E. intestinalis*, we used FoldSeek^56^ to search the AlphaFold Database^57^ of predicted protein structures on March 30, 2023. We initiated the search using the predicted structure of Eint_081460 as a probe and restricted the search to the suborder Apansporoblastina, which includes *E. intestinalis*. This resulted in 196 hits from a number of different species; these sequences were each used to initiate a BLASTp restricted to *E. intestinalis* ATCC 50506. The proteins identified from all of these searches were combined in a single list and duplicates were removed, resulting in 14 unique *E. intestinalis* proteins. Re-predicting these structures of these 14 proteins through ColabFold^58^ strongly suggested the presence of ricin B domains in 12 of 14 proteins, based upon manual examination of the predicted folds and consistency in the structure prediction across the 5 top ranked predictions. For the 13th protein (Eint_070940), the predictions were more variable, with only one of the top five predictions resembling a ricin B domain. This initial prediction included the N-terminal signal peptide and a predicted disordered region from the C-terminus, which can sometimes interfere with AlphaFold2 predictions of neighboring regions. After deleting these regions, the remaining sequence (residues 16-156) was predicted to adopt a ricin B domain fold with good agreement across the top 5 predictions. For the 14th and final protein (Eint_071030), the predictions had lower pLDDT scores and were more variable between predictions, suggesting a less reliable prediction. However, the best scoring predictions resembled an RBL fold, with similar topology and connectivity and could be superposed on other RBL domains. These predictions deviated primarily in the prediction of first and final β-strands, and similar results were obtained with the Eint_071030 ortholog from *E. cuniculi* (ECU07_1070), which shares ∼70% sequence identity. Thus, it appears that *E. intestinalis* likely encodes 14 ricin B domain-containing proteins in its genome (Extended Data Fig. 5).

### Prediction of secreted proteins

The sequences of all annotated proteins from the *E. intestinalis* ATCC 50506 genome (a total of 1,934 sequences) were downloaded from Uniprot on April 6, 2023. Prediction of possible signal peptides was carried out using SignalP-6.0^59^. However, only a relatively small fraction of the *E. intestinalis* proteome was predicted to be secreted: only 3.4% of the proteome (66 proteins) was predicted to have a signal peptide with a probability of 0.5 or higher. In contrast, a similar analysis of the Plasmodium falciparum genome predicted that 8.8% of the proteome had a signal peptide with a probability of 0.5 or higher. Several proteins expected to be secreted from *E. intestinalis* had much lower signal peptide probabilities, including PTP3 (probability = 0.34) and several Ricin B domain-containing proteins (e.g., Eint_081460, probability = 0.16). Microsporidia are evolutionarily divergent, and protein secretion has primarily been studied in model organisms. Thus, we hypothesized that prediction tools may still detect many signal peptides in *E. intestinalis* proteins, but that *E. intestinalis* signal peptides may result in lower scores due to lineage specific differences in signal peptide lengths, composition, cleavage site preferences, etc. Indeed, examination of a waterfall plot of predicted signal peptide probabilities for all *E. intestinalis* proteins indicates that, as expected, most proteins are predicted to not have signal peptides (Extended Data Fig. 6). A distinct dogleg separates proteins with very low probabilities of a signal peptide from proteins with a probability greater than ∼0.05. Thus, we predict a protein to be secreted if it has a probability of a signal peptide >0.05 according to SignalP-6.0, which includes most of the proteins that we infer are likely to be secreted based on their functional annotations and predicted structures, while including a relatively small number of proteins that are unlikely to have true signal peptides (e.g., Eint_010070, an ABC transporter, where the first transmembrane helix appears to be mistaken for a signal peptide with a probability of 0.11).

### Secretion Assay

To assess whether the N-terminal region of an *E. intestinalis* protein can function as a signal peptide and direct the protein to the secretory pathway, we fused codons for the N-terminal 40 amino acids of the protein to the N-terminus of sfGFP, and cloned the resulting open reading frame into a mammalian expression vector, pBEL1784 (a derivative of pcDNA3.4). Briefly, sfGFP was amplified from template DNA, with the additional 40 codons from the *E. intestinalis* proteins encoded by the primer/Ultramer (IDT). The resulting PCR products were cloned via Gibson assembly^60^ in pBEL1784 previously digested with EcoRI-HF and NotI-HF (New England Biolabs). In addition, a sfGFP secreted positive control (rabbit IgH signal peptide) and a non-secreted/cytoplasmic sfGFP negative control (no signal peptide) were similarly constructed. The resulting plasmids, including the putative signal peptide sequences, are listed in Supplementary Table 7.

Prior to transfection, Expi293F cells (Gibco A14635) were grown in a 250 mL polycarbonate shaker flask using Expi293 Expression Medium (Gibco) at 37°C and 125 rpm, and with 8% CO_2_. Transfection of the plasmids listed in Supplementary Table 7 into Expi293F cells was performed according to the manufacturer’s instructions using the Gibco™ ExpiFectamine™ 293 Transfection Kit (Gibco™ A14524). Following transfection, the cells were grown in a sealed 24-well deep well plate (AeraSeal BS25; Axygen™ PDW10ML24CS) at 37°C and 235 rpm without supplemental CO_2_. 24 hours post-transfection, 100 μL of cells were collected from each well and centrifuged at 1,000 g for 5 minutes at room temperature. Samples of supernatant were carefully removed without disturbing the cell pellet. The remaining cell pellet was resuspended in 100 μL of Dulbecco’s Phosphate-Buffered Saline (DPBS) (Gibco). SDS sample loading buffer was added to the supernatant samples and resuspended cells to a final concentration of 1X, then heated at 100°C for 5 minutes. The resulting whole cell lysate samples were mechanically disrupted using a G-27 needle after boiling the sample, to shear genomic DNA.

Western blots were performed on the supernatant samples to assess secretion of GFP into the culture medium, and on the whole cell lysates to assess cellular expression levels for each construct. All samples were run on 4– 20% Criterion TGX Stain-Free Protein Gels (Bio-Rad Laboratories). After electrophoresis, proteins were transferred to a nitrocellulose membrane (Bio-Rad Laboratories) using the Trans-Blot Turbo Transfer System (Bio-Rad Laboratories). Membranes were blocked with 1X TBS (150 mM NaCl, 20 mM Tris; pH 7.6) + 0.1%Tween 20 (TBST) + 5% milk for an hour at room temperature and then washed twice with TBST for five minutes. The membranes were incubated with primary antibodies against GAPDH (mouse anti-GAPDH antibody at 0.5 μg/mL (abcam ab125247)) and GFP (custom rabbit polyclonal anti-GFP antibody at 0.75 μg/mL (provided by the Foley laboratory)) in TBST + 5% BSA for one hour at room temperature. The membranes were washed with TBST three times for five minutes and were incubated with goat anti-mouse IgG polyclonal antibody (IRDye 680RD, LI-COR Biosciences 926-68070) at 0.1 μg/mL and goat anti-rabbit IgG polyclonal antibody (IRDye 800CW, LI-COR Biosciences 925-32211) at 0.1 μg/mL in TBST + 5% milk for an hour at room temperature. A final wash was done with TBS three times for ten minutes. The membranes were then imaged using the LI-COR Odyssey XF (LI-COR Biosciences).

## Data Availability

All sequencing data have been deposited in NCBI’s Gene Expression Omnibus under accession no. GSE268707.

## Code Availability

All codes used to perform the analysis and reproduce the figures are available on GitHub: https://github.com/kaciemccarty/E.-intestinalis-scRNAseq.git

## Acknowledgements

We thank Noelle Antao, Fred Rubino, Alice Herneisen, and Meike Dittmann for feedback on our manuscript and all members of the Bhabha+Ekiert labs for helpful discussions. We thank Victor Torres for providing PBMCs; Adriana Heguy and Peter Meyn from the NYU Genome Technology Center for the single cell library preparation and sequencing; Michael Cammer from the NYU Microscopy Laboratory for discussion of light microscopy experiments. We gratefully acknowledge the following funding sources: SSP-2018-2737 (Searle Scholars Program, to G.B.); R01AI147131 (NIAID, to G.B.); Irma T. Hirschl Career Scientist Award (to G.B.); and the Thailand Science Research and Innovation Fund, Chulalongkorn University, Grant no. 4709739 (to P.J.). G.B. is a Pew Scholar in the Biomedical Sciences, supported by The Pew Charitable Trusts (PEW-00033055).

