## Extended Data and Supplementary Information for "scRNA-seq reveals transcriptional dynamics of *Encephalitozoon intestinalis* parasites in human macrophages"

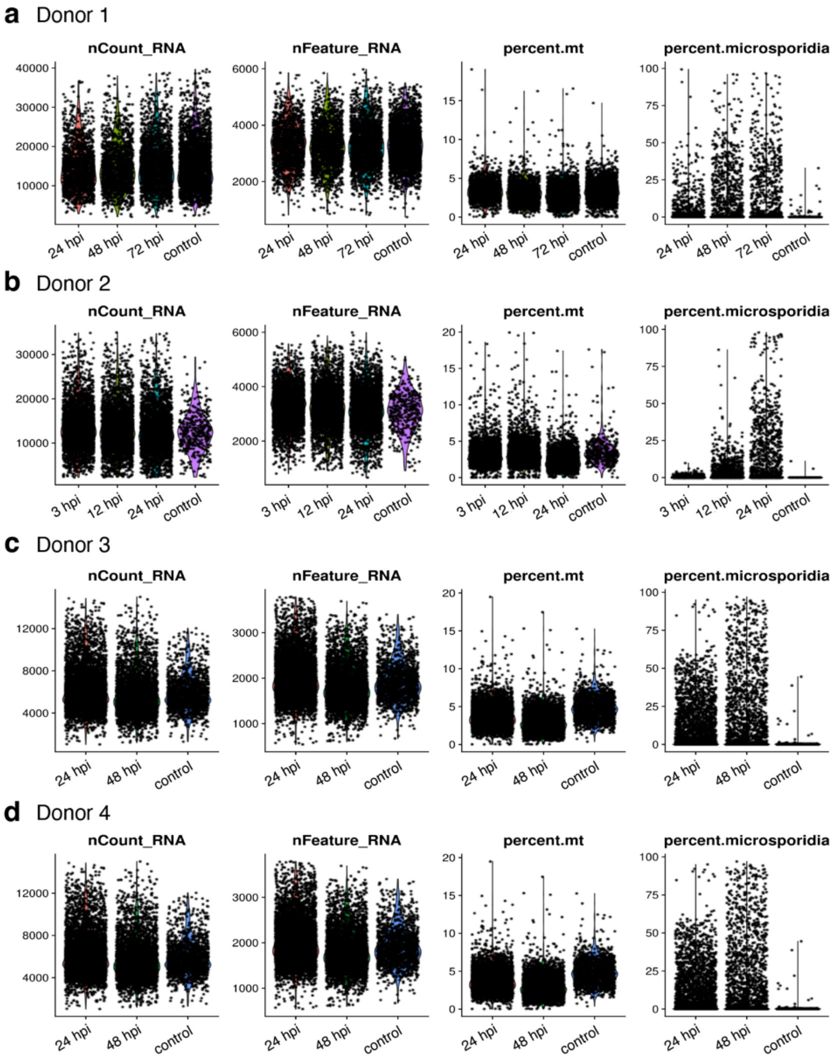

**Extended Data Figure 1. Quantification of RNA transcripts and genes.**
**A-D.** Quantification of transcripts (nCount), genes (nFeature), percent mitochondria (percent.mt), and percent microsporidia in each cell from each donor from the Total transcriptome.

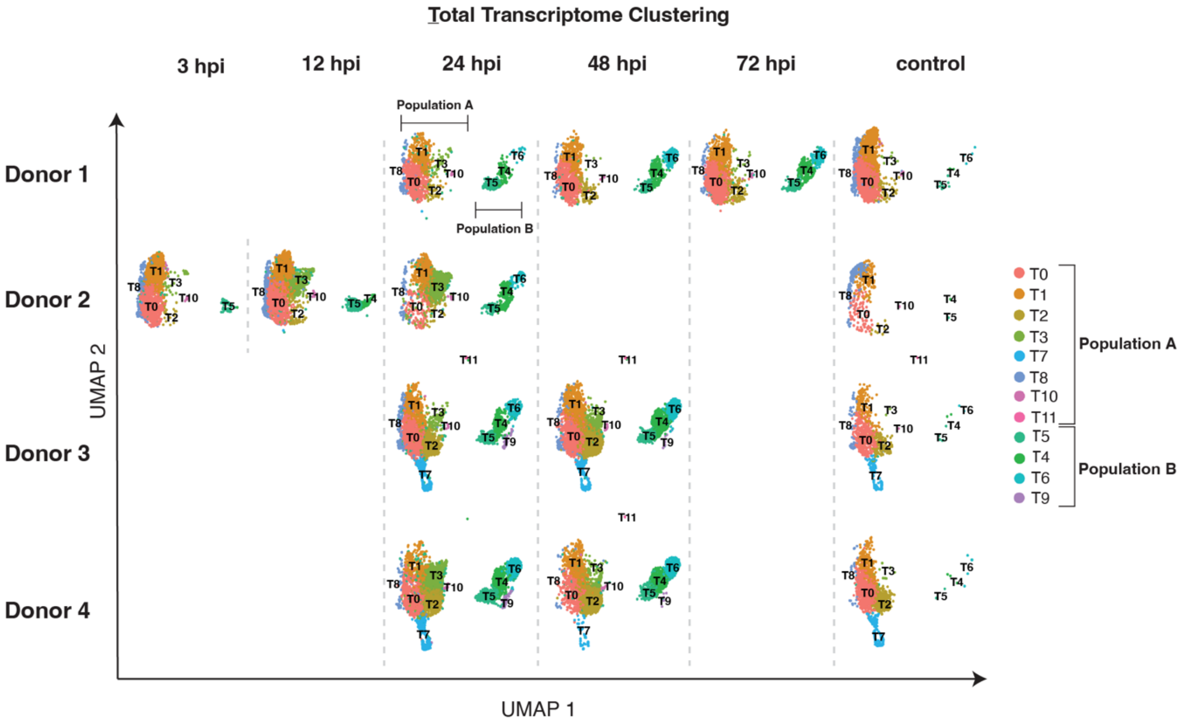

**Extended Data Figure 2. Total transcriptome analysis.**
UMAP plot separated by time point and donor.

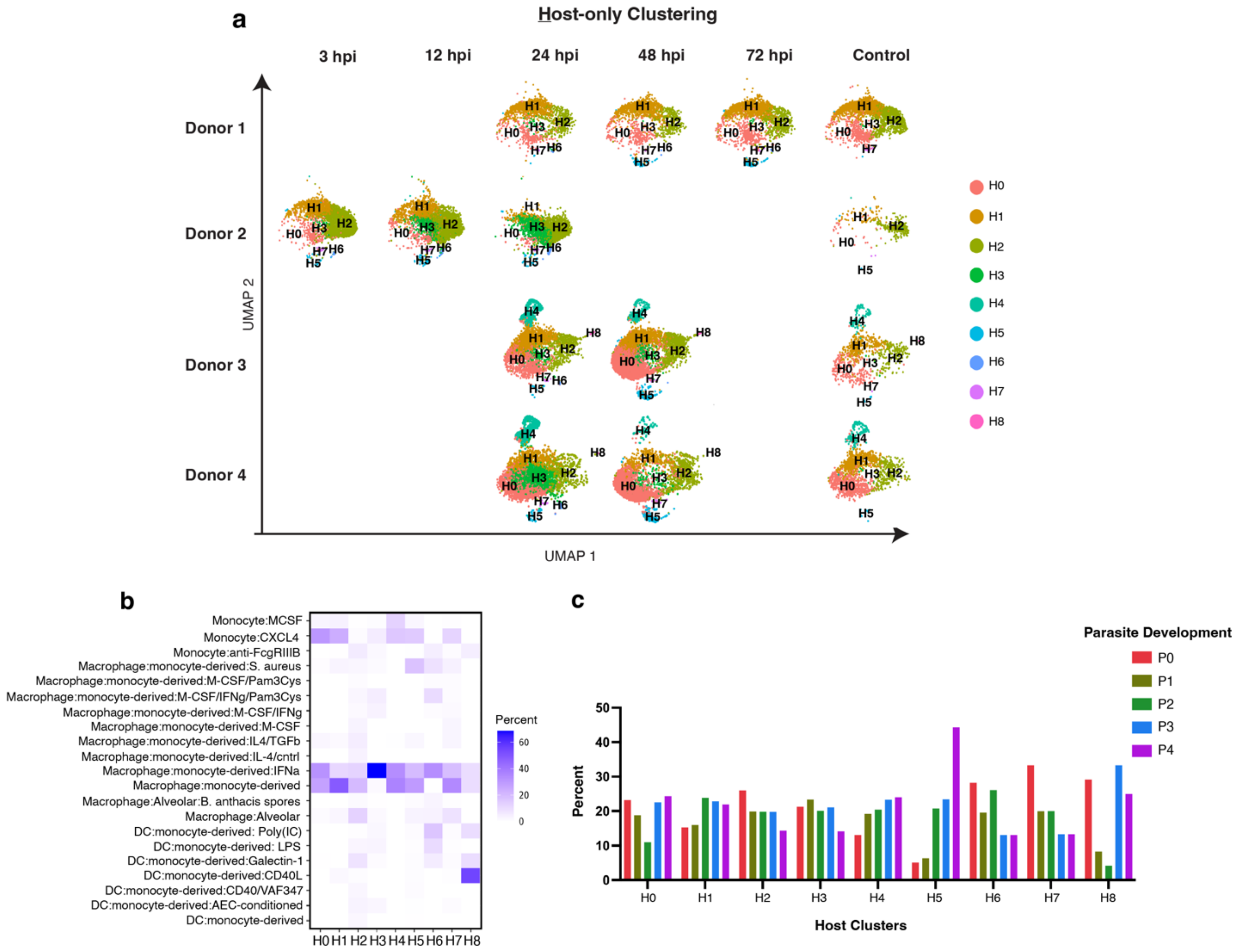

**Extended Data Figure 3. Human-only transcriptome analysis.**
**A.** UMAP plot separated by time point and donor. **B.** Heatmap of the percentage of cells from each cluster annotated as a specific cell type based on the Human Primary Cell Atlas (HPCA)**. C.** Quantification of the percentage of parasite developmental stages found within the human-only clusters.

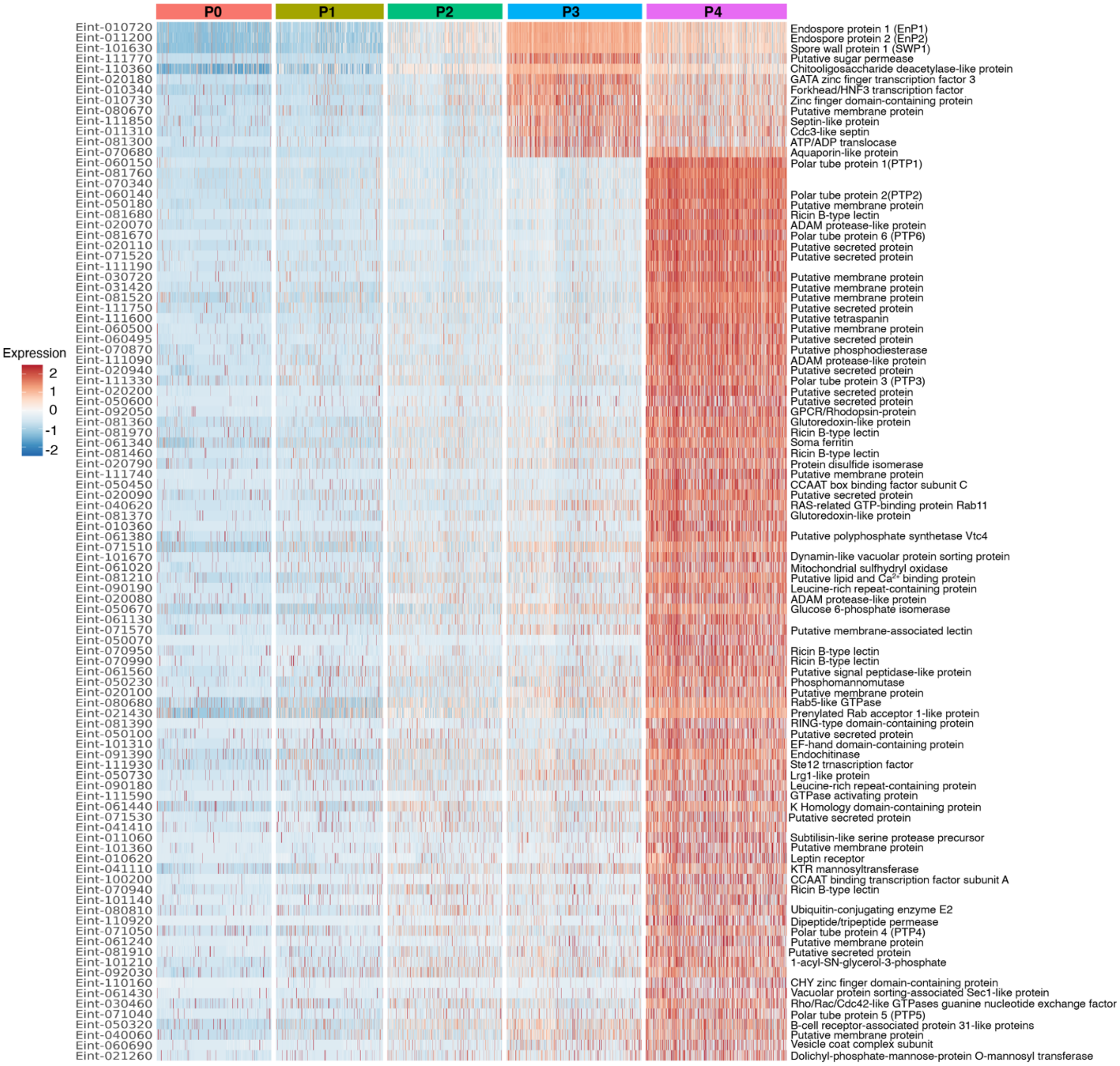

**Extended Data Figure 4. Differential gene expression analysis of the parasite-only transcriptome.**
Heatmap of differentially expressed genes with a fold change of ≥ 2.0.

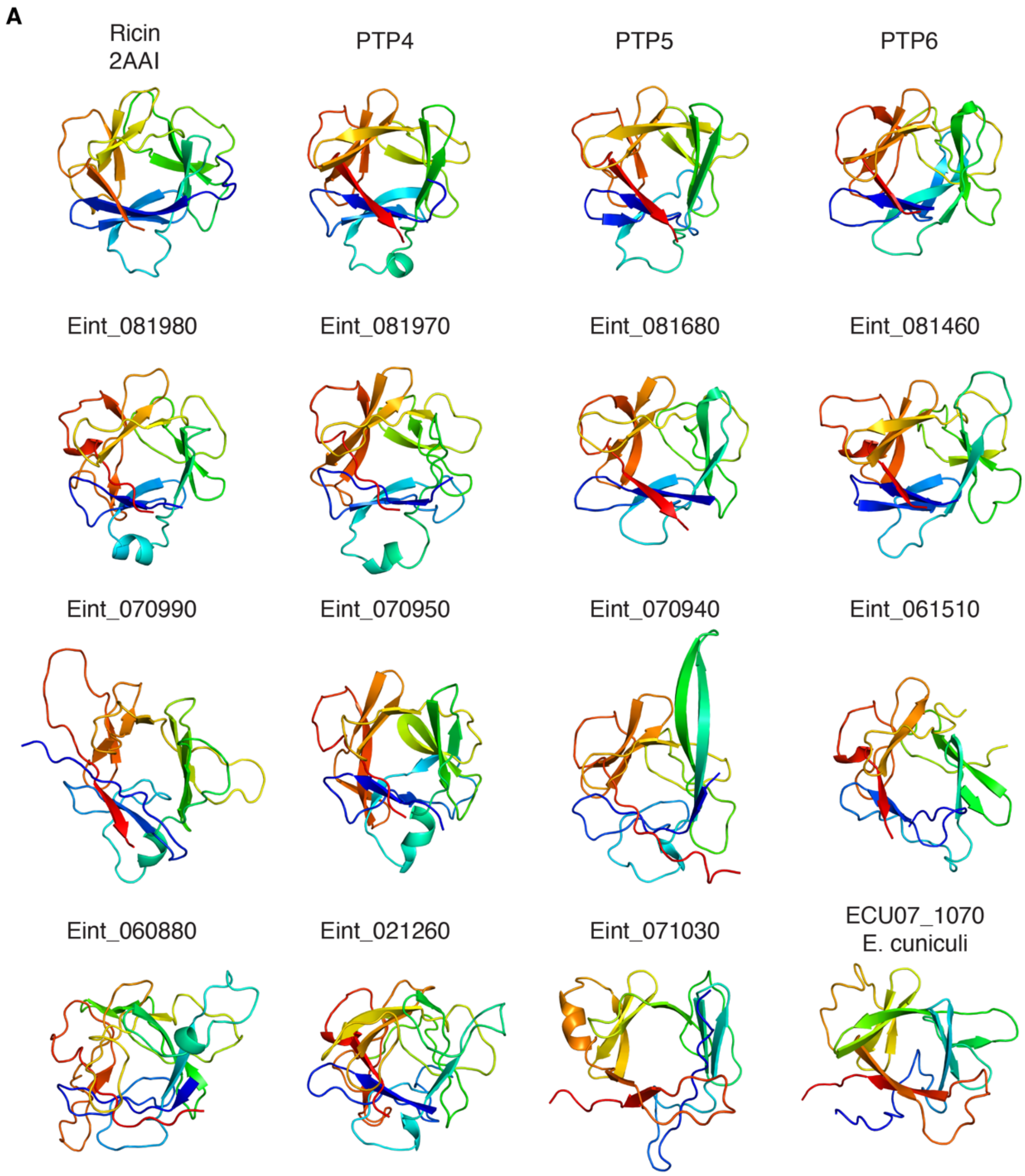

**Figure legend on next page.**

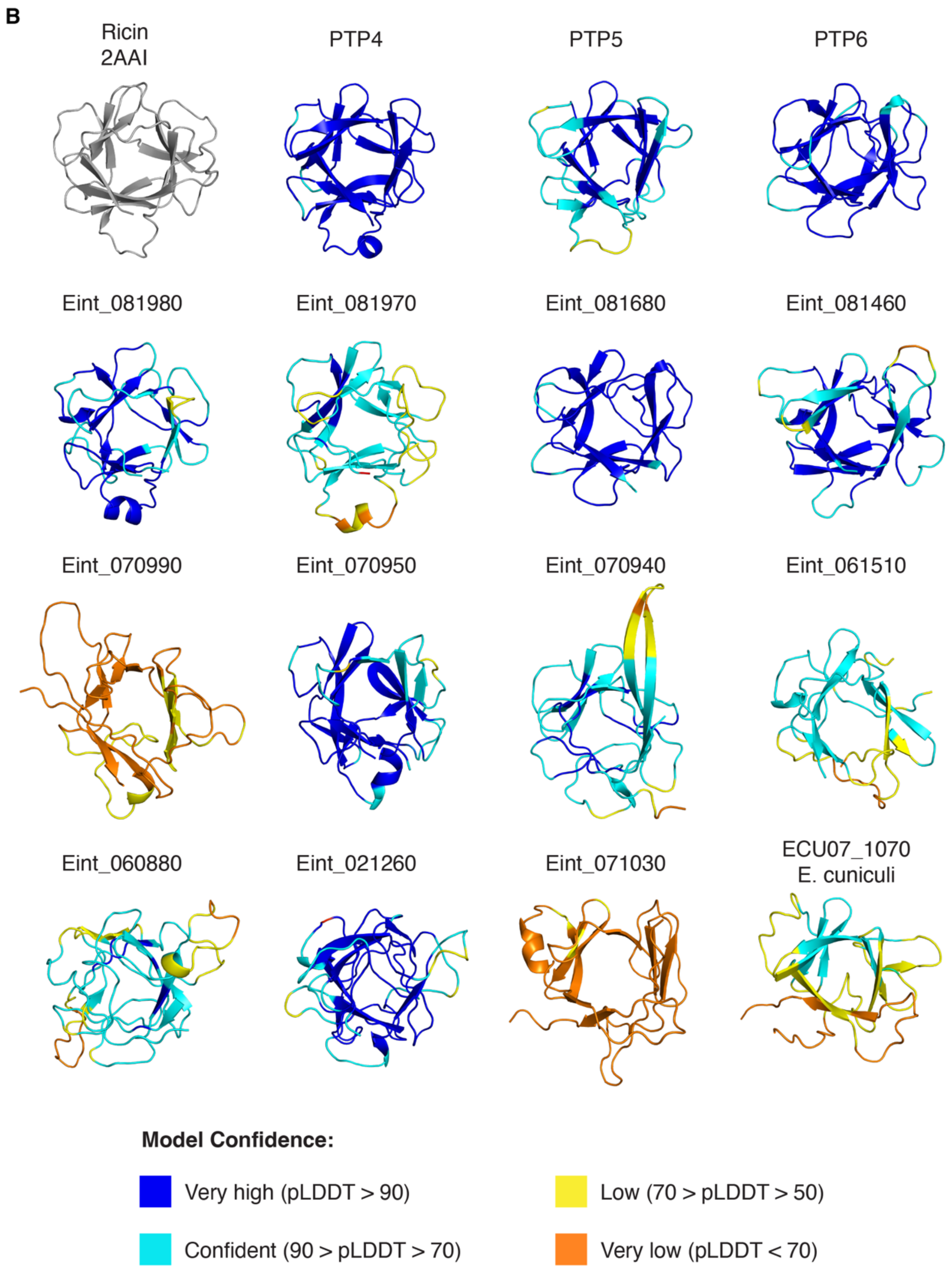

**Extended Data Figure 5. Alpha fold prediction of Ricin B-type lectin domain containing proteins**

**A.** Alpha fold predictions of putative Ricin B-type lectin domain containing proteins from *E. intestinalis* and *E. cuniculi* (ECU07_1070) compared to the Ricin toxin from *Ricinus communis* (PDB: 2AAI). **B.** Alpha fold predictions colored by pLDDT.

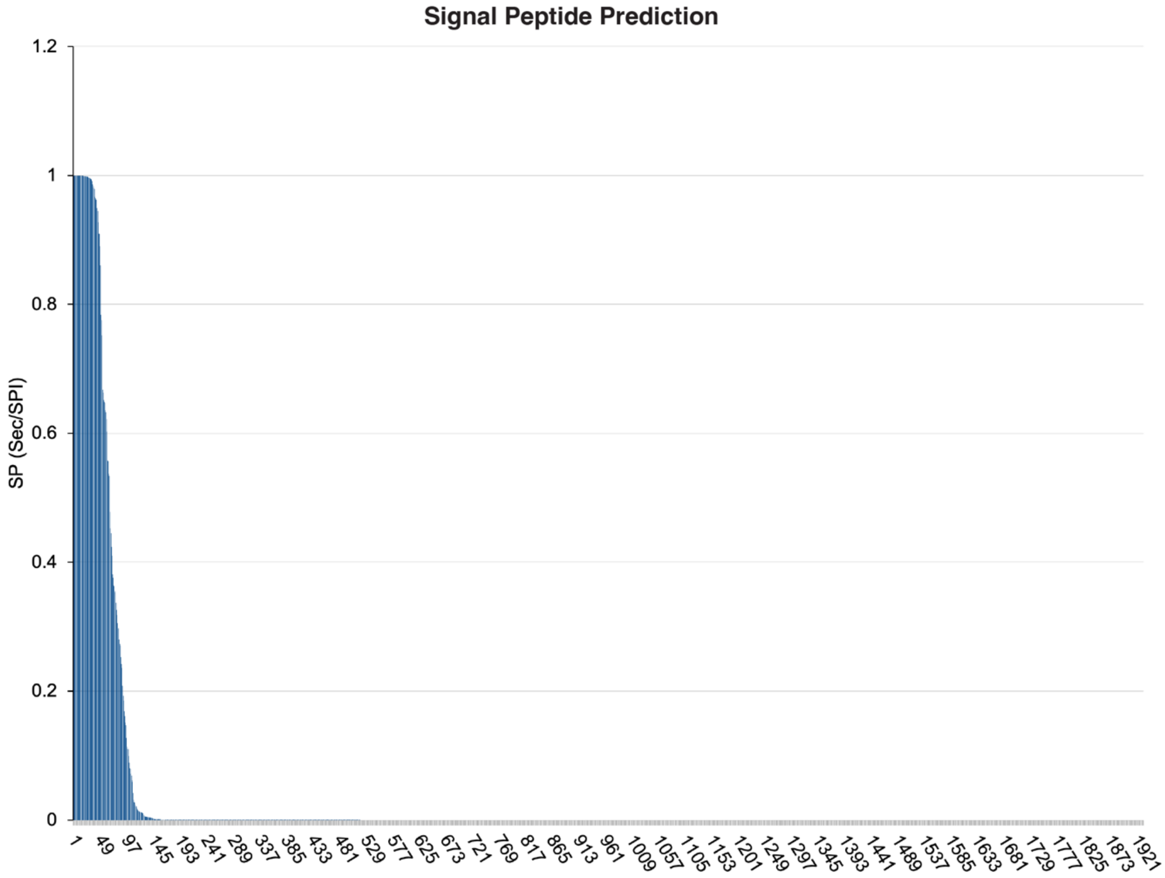

**Extended Data Figure 6. Prediction of secreted proteins**
Waterfall plot representing the number of proteins predicted to contain a signal peptide.

**Supplementary Data 1.** Average expression of parasite genes across all clusters.

- See 'Supplementary_Data_1.xls'

**Supplementary Data 2.** Differentially expressed parasite genes from each cluster. Note that the lowest p-values that R could show is 2.23e-308. Lower p-values than this will be indicated as zero.

- See 'Supplementary_Data_2.xls'

**Supplementary Table 1.** Number of cells that were included in the analyses after filtering out bad quality cells.

| Timepoint | Donor 1 | Donor 2 | Donor 3 | Donor 4 | Total |
|---|---|---|---|---|---|
| **Control** | 2,842 | 397 | 825 | 1,549 | 5,613 |
| **3 hpi** | - | 3,283 | - | - | 3,283 |
| **12 hpi** | - | 3,281 | - | - | 3,281 |
| **24 hpi** | 1,536 | 3,101 | 3,465 | 5,513 | 13,615 |
| **48 hpi** | 1,604 | - | 5,964 | 4,406 | 11,974 |
| **72 hpi** | 2,115 | - | - | - | 2,115 |
| **Summary** | 8,097 | 10,062 | 10,254 | 11,468 | 39,881 |

**Supplementary Table 2.** Number of infected cells found in each donor.

| Timepoint | Donor 1 | Donor 2 | Donor 3 | Donor 4 | Total |
|---|---|---|---|---|---|
| **Control** | 17 | 2 | 10 | 10 | 39 |
| **3 hpi** | - | 45 | - | - | 45 |
| **12 hpi** | - | 479 | - | - | 479 |
| **24 hpi** | 312 | 631 | 775 | 1524 | 3242 |
| **48 hpi** | 517 | - | 1188 | 1087 | 2792 |
| **72 hpi** | 459 | - | - | - | 459 |
| **Summary** | 1025 | 1157 | 1973 | 2621 | 6776 |

**Supplementary Table 3.** Number of cells found in each donor at each timepoint in each cluster from the "total" transcriptome dataset.

- See 'Extended_Data_Table_3.xls'

**Supplementary Table 4.** Differential gene expression analysis showing the top thirty highly expressed genes in each cluster.

- See 'Supplementary_Table_4.xls'

**Supplementary Table 5.** *E. intestinalis* proteins predicted to contain a signal peptide using SignalP6.0.

- See 'Supplementary_Table _2.xls'

**Supplementary Table 6.** Number of cells that were used for scRNA-seq. "Double" is used to annotate cells containing more than one bound hashing antibody. "Negative" is used to annotate that there is no hashing antibody bound to the cells.

| Timepoint | Donor 1 | Donor 2 | Donor 3 | Donor 4 | Total |
|---|---|---|---|---|---|
| **Control** | 2,870 | 415 | 830 | 1,572 | 5,687 |
| **3 hpi** | - | 3,434 | - | - | 3,434 |
| **12 hpi** | - | 3,483 | - | - | 3,483 |
| **24 hpi** | 1,559 | 3,219 | 3,500 | 5,583 | 13,861 |
| **48 hpi** | 1,631 | - | 5,984 | 4,431 | 12,046 |
| **72 hpi** | 2,136 | - | - | - | 2,136 |
| **Double** | 878 | 628 | 6,535 | | 8,041 |
| **Negative** | 1,026 | 1,100 | 1,920 | | 4,046 |
| **Summary** | 10,100 | 12,279 | 30,355 | | 52,734 |

**Supplementary Table 7.** Plasmids used in this study

| Plasmid ID | Putative Signal Peptide Source | Putative Signal Peptide Sequence | Plasmid Source | Addgene ID |
|---|---|---|---|---|
| pBEL2837 | Rabbit IgH (positive control) | METGLRWLLLVAVLKGVQC | This study | |
| pBEL2838 | No Signal Peptide (cytoplasmic) | | This study | |
| pBEL2957 | Eint_060150 | MKGISKVLSASIVLMKLKGVYSTTVLCGDSTQGLQGTTQP | This study | |
| pBEL2999 | Eint-060140 | MLLLLSAVAFVSATAVQSGVVSQPTTPIPILPGQPMGGMA | This study | |
| pBEL3000 | Eint_111330 | MLLFLLCLYNMEVRSETKVSPDAIHRSKEANTGKGMDYGN | This study | |
| pBEL3001 | Eint_071050 | MEPGLILMFMSIFVSARDRELEEFVEKDIKVFFSSYPTQI | This study | |
| pBEL3002 | Eint_071040 | MILILVLGMAFSYPLYLDSFTNKPIRILAAEYHLLVSELK | This study | |
| pBEL3003 | Eint_081670 | MKSIFVVFQLVKVISGFTIFTKDGQRKHLVKGGIWRSPDY | This study | |
| pBEL3004 | Eint_081980 | MKLTIVLMISFAFCISDPIK | This study | |

| | | | |
|---|---|---|---|
| | | VKIVSKVDPKKYLSLFSG EV | |
| pBEL3005 | Eint_081970 | MRLLVFCMTSLVMGSRM VRITPRTHPDFVVAIVPPK LGGK | This study |
| pBEL3006 | Eint_081680 | MKVISFGCIISTAFAFMIT SENDEKKFVSNSSGKAV MTTT | This study |
| pBEL3007 | Eint_081460 | MLFIYLALTYAMEEMGYI FSEGKQRFLTRNGVNIVV GRSK | This study |
| pBEL3008 | Eint_071030 | MEFIFLLLLSGACGHIKNP VEIRLFLKPDLELKVGRIP LI | This study |
| pBEL3009 | Eint_070990 | MNVYILLHLGMVVSRMGI AIKSMMSGRYLNLRTLRL EPVD | This study |
| pBEL3010 | Eint_070950 | MLLLFFAFAYSFQRGYG PCGRSNRHLRNSDYDGV MMLNVL | This study |
| pBEL3011 | Eint_070940 | MKRRAYICFFLGIATCEIF LTTIRPIDNPNVLMVSGK NVV | This study |
| pBEL3012 | Eint_061510 | MISMLVFLPFVICDRLLRL KHKDSQYVTNNNGILKLD FPL | This study |
